# Co-existing feedback loops generate tissue-specific circadian rhythms

**DOI:** 10.1101/304451

**Authors:** J. Patrick Pett, Matthew Kondoff, Grigory Bordyugov, Achim Kramer, Hanspeter Herzel

## Abstract

Gene regulatory feedback loops generate autonomous circadian rhythms in mammalian tissues. The well-studied core clock network contains many negative and positive regulations. Multiple feedback loops have been discussed as primary rhythm generators but the design principles of the core clock and differences between tissues are still under debate.

Here we use global optimization techniques to fit mathematical models to circadian gene expression profiles for different mammalian tissues. It turns out that for every investigated tissue multiple model parameter sets reproduce the experimental data. We extract for all model versions the most essential feedback loops and find auto-inhibitions of Period and Cryptochrome genes, *Bmal1/Rev-erb-α* loops, and repressilator motifs as possible rhythm generators. Interestingly, the essential feedback loops differ between tissues, pointing to specific design principles within the hierarchy of mammalian tissue clocks. Self-inhibitions of *Per* and *Cry* genes are characteristic for models of SCN clocks, while in liver models many loops act in synergy and are connected by a repressilator motif. Tissue-specific use of a network of co-existing synergistic feedback loops could account for functional differences between organs.

## 1 Introduction

Many organisms have evolved a circadian (∼24 hour) clock to adapt to the 24 h period of the day/night cycle [1]. In mammals, physiological and behavioral processes show circadian regulation including sleep-wake cycles, cardiac function, renal function, digestion, and detoxification [2]. In most tissues, about 10% of genes have circadian patterns of expression [3, 4]. Surprisingly, the rhythmicity of clock-controlled genes is highly tissue specific [4–6].

Circadian rhythms are generated in a cell-autonomous manner by transcriptional/translational feedback loops [7] and can be monitored even in individual neurons [8] or fibroblasts [9].

Ukai and Ueda [10] depict the mammalian core clock as a network of 20 transcriptional regulators (10 activators and 10 inhibitors) acting via enhancer elements in their promoters such as E-boxes, D-boxes and ROR-elements. Since many of these regulators have similar phases of expression and DNA binding [11, 12], the complex gene regulatory network has been reduced by Korencic et al. [6] to just 5 regulators representing groups of genes: the activators *Bmal1* and *Dbp* and the inhibitors *Per2, Cry1* and *Rev-Erba* (see Figure 1A).

**Figure 1:**
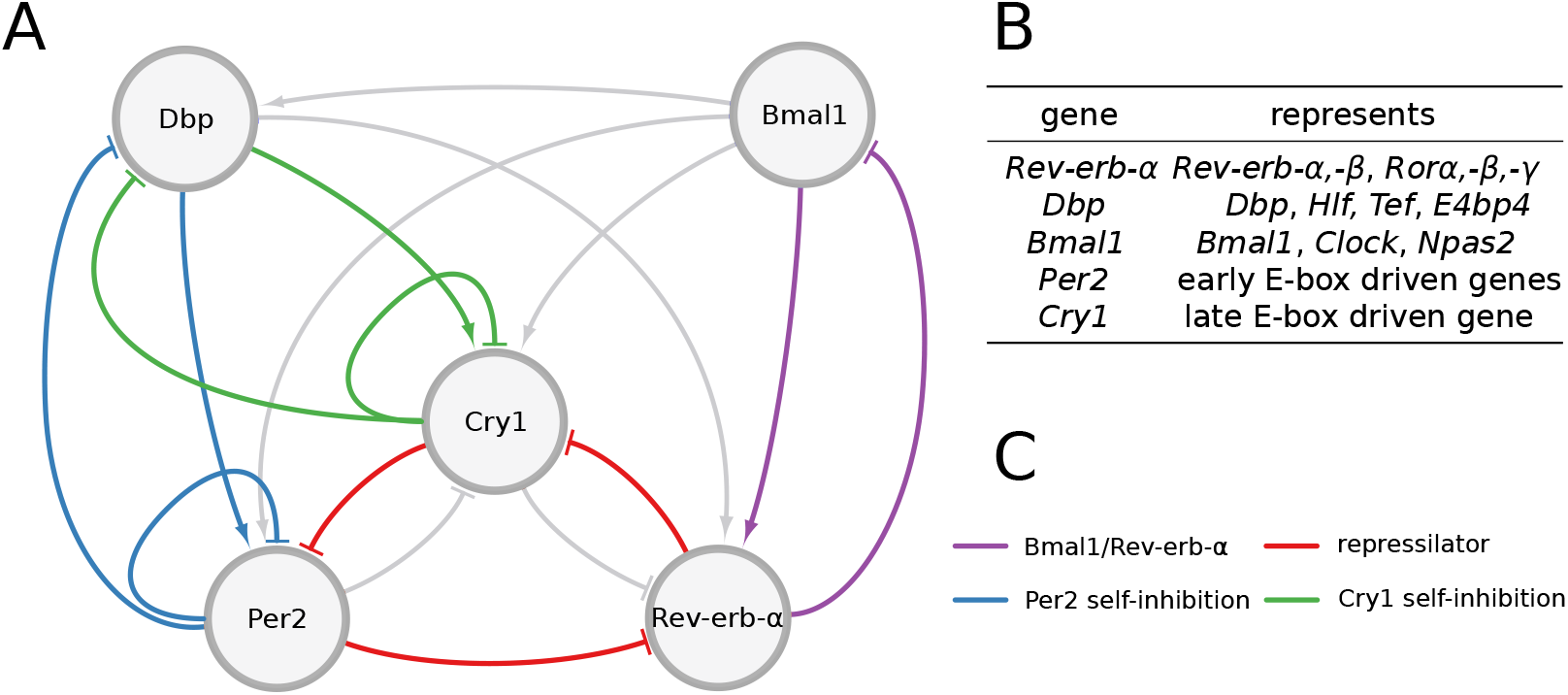
Network of the core clock model. The graph comprises 7 activations and 10 inhibitions forming several negative feedback loops. Four loops that are mainly discussed in the literature and were most often found by our analysis are marked in different colors. Note, that for *Per2* and *Cry1* auto-inhibitions also extensions via the gene *Dbp* are counted.

Even this condensed network contains 17 regulations constituting multiple negative and positive feedback loops [13]. In order to generate self-sustained oscillations negative feedback loops are essential [14, 15]. Originally, the self-inhibitions of Period and Cryptochrome genes have been considered as the primary negative feedback loops [16]. Later, computational modeling [17] and double-knockout experiments suggested that also the *Rev-Erb* genes play a dominant role in rhythm generation [18]. Recently, it has been shown that also a combination of 3 inhibitors forming a repressilator [19] can reproduce expression patterns in liver, adrenal gland and kidney [13].

Despite many experimental and theoretical studies major questions remain open: What are the most essential feedback loops in the core clock network? Do dominant loop structures vary across tissues?

Here, we use global optimization techniques to fit our 5-gene model to expression profiles in different mammalian tissues (adrenal gland, kidney, liver, heart, skeletal muscle, lung, brown adipose, white adipose, SCN and cerebellum) [3]. We find that for any given tissue multiple parameter sets reproduce the data within the experimental uncertainties. By clamping genes and regulations at non-oscillatory levels [13] we unravel the underlying essential feedback loops in all these models. We find auto-inhibitions of Period and Cryptochrome genes, *Bmal1/Rev-erb-α* loops and repressor motifs as rhythm generators. The role of these loops varies between organs. For example, in liver repressilators dominate whereas *Bmal1/Rev-erb-α* loops are found in the heart. Clustering of the model parameter sets reveals tissue-specific loop structures. For example, we rarely find the repressilator motif in brain, heart and muscle tissues due to earlier phases and small amplitudes of *Cry1*. We discuss that the co-existence of functional feedback loops increases robustness and flexibility of the circadian core clock.

## 2 Results

### 2.1 A 5-gene regulatory network represents most essential loops

Here, we derive a gene regulatory network that can be fitted successfully to available transcriptome, proteome and ChIP-seq data. We will use the model to explore tissue-specific regulations.

Many circadian gene expression profiles for mouse tissues are available [5, 20, 21]. Here we focus on data sets from tissues spanning 48 hours with a two hour sampling [3]. These comprehensive expression profiles are particularly well suited to study tissue differences. For mouse liver also proteome data [22] and ChIP-seq data [11, 23, 24] are available with lower resolution.

Using global parameter optimization we fit tissue-specific model parameters directly to the gene expression profiles of Zhang et al. [3]. Proteome and ChIP-seq data are primarily used to specify reasonable ranges of the delays between transcription and the action of activators and repressors. The ranges of degradation rates have been adapted to large-scale studies measuring half-lifes of mRNAs [25, 26] and proteins [27].

Quantitative details of activation and inhibition kinetics are not known due to the high complexity of transcriptional regulation. The transcriptional regulators are parts of MDa complexes [28] including HATs and HDACs [29]. Details of DNA binding, recruitment of co-regulators and histone modifications are not available [30]. Thus we use heuristic expressions from biophysics [31] to model activation and inhibition kinetics. Exponents represent the number of experimentally verified binding sites [32] (see Supplement S1) and the parameters were assumed to be in the range of the working points of regulation.

In order to justify the topology of our reduced gene regulatory network we analyze the amplitudes and activation phases of all the 20 regulators described in Ukai and Ueda [10] (Figure 2). Repressor phases were inverted by 12 h to reflect the maximal activity and allowing direct comparison with activators.

**Figure 2:**
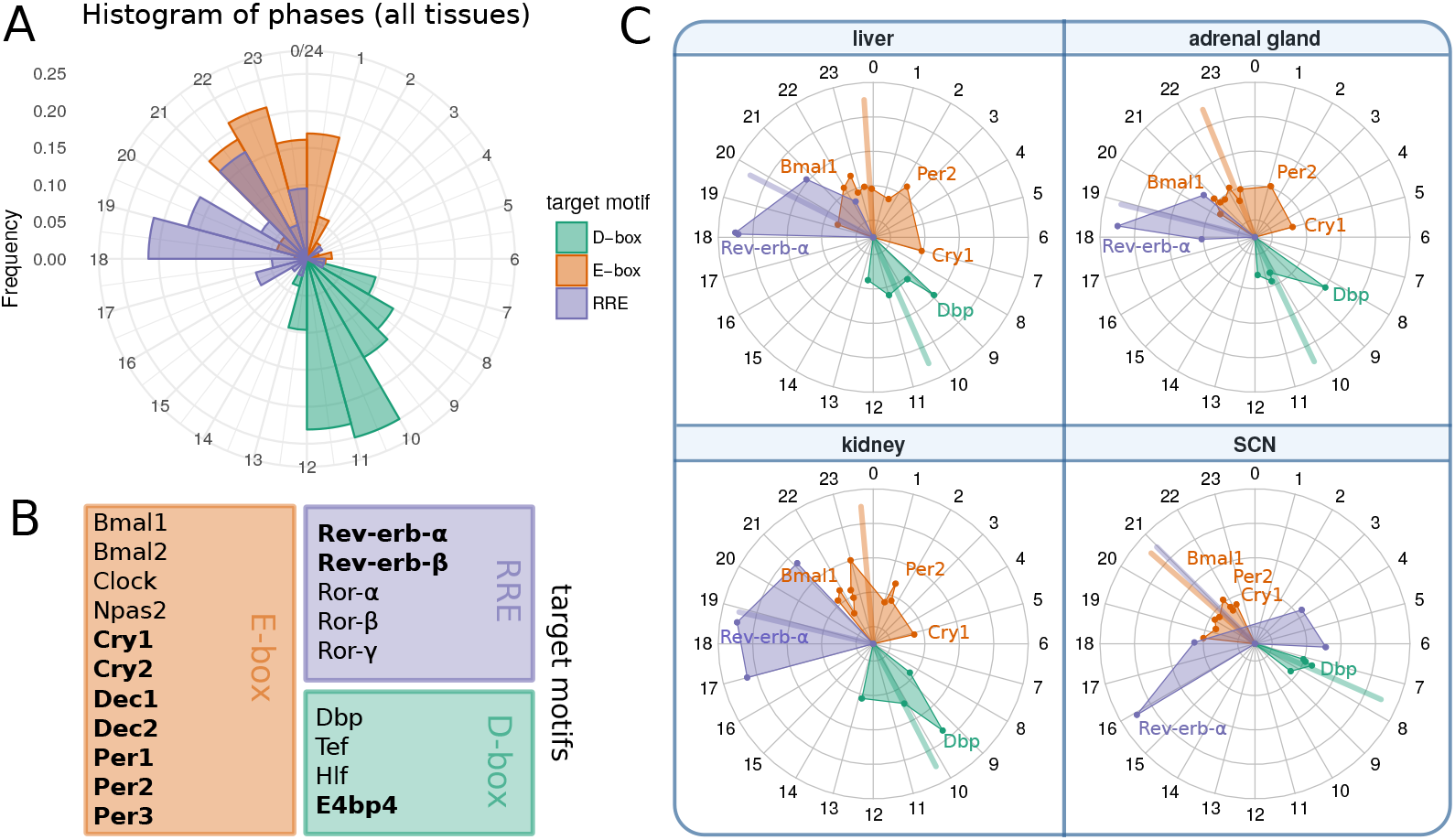
Circular plots of 20 regulators reveal redundancies and serial inhibition [10, 12]. They represent peak phases of mRNA expression in multiple tissues [3]. Note that repressor phases were inverted by 12 h to allow direct comparison with activators. (A) Histogram of the phase distribution over all tissues. (B) List of genes represented in circular plots and their corresponding target motifs. Repressors are marked in bold. (C) Phases of core clock genes in selected tissues. Colored lines correspond to the circular mean of the respective groups. Amplitudes are linearly scaled. The differences between SCN and other tissues are particularly notable (for example the earlier *Cry1* peak). Antagonistic regulations of *Rev-erb* and *Ror* in the SCN can be modeled by reduced inhibition strength of *Rev-erb-α*.

Figures 2 shows that the 5 genes binding to ROR-elements (RREs) and the 4 genes binding to D-boxes cluster at specific phases. Consequently, we represent these regulators by selected genes with large amplitudes: *Rev-erb-α* and *Dbp*. Since the other RRE and D-box regulators peak at similar or directly opposed phases their additional regulation can be taken into account by the fitting of activation and inhibition parameters.

The regulation via E-boxes is quite complex [11, 30, 33, 34]. In addition to the activators *Bmal1* and *Bmal2* we have their dimerization partners *Clock* and *Npas2* and their competitors *Dec1* and *Dec2*. Furthermore, there are the early E-box targets *Per1, Per2, Per3* and *Cry2* and the late gene *Cry1*. We model this complicated modulation by 3 representative genes: *Bmal1* as the main activator and *Per2* and *Cry1* as early and late E-box target.

In summary, the reduced gene regulatory network consists of 5 genes and 17 regulations (see Figure 1). All regulations and the number of binding sites have been confirmed by several experimental studies discussed in detail in [32]. Interestingly, liver proteomics [22] and ChIP-seq data are consistent with morning activation via *Bmal1*, evening activation by *Dbp* and sequential inhibition by *Rev-erb-α, Per2*, and *Cry1*. Recent detailed biochemical experiments support the notion that there are distinct inhibition mechanisms associated with *Per2* and *Cry1* [30]. The essential role of late *Cry1* inhibition has been stressed also by Ukai et al. [35] and Edwards et al. [36].

As discussed above our model is fitted directly to mRNA time series collected for different tissues at 2 h intervals for a total duration of 2 days. The transcriptional/translational loops are closed by delayed activation or repression realized by the corresponding proteins. Since most quantitative details of posttranscriptional modifications, complex formations, nuclear import and epigenetic regulations are not known, we simplify all these intermediate processes by using explicit delays. Thus we describe the core clock network by5 delay-differential equations (DDEs) with 34 kinetic parameters (see Supplement 1 for the complete set of equations).

The model constitutes a strongly reduced network that approximates the highly complex protein dynamics by delays. Inhibition strengths are represented by a single parameter whereas modeling of activation requires 2 parameters: maximum activation and threshold levels. While keeping in mind the proposed simplifications, the resulting tissue-specific models can still be regarded as a biologically plausible regressions of the underlying biological dynamics.

In Figure 3A we show examples of simulations fitted to the corresponding gene expression patterns. After successful parameter optimization as described below the differences between data and models are comparable to experimental uncertainties quantified by comparing different studies [3, 21, 37] and by studying differences between the first and second day of the expression profiles by Zhang et al. [3] (see Supplement S3).

**Figure 3:**
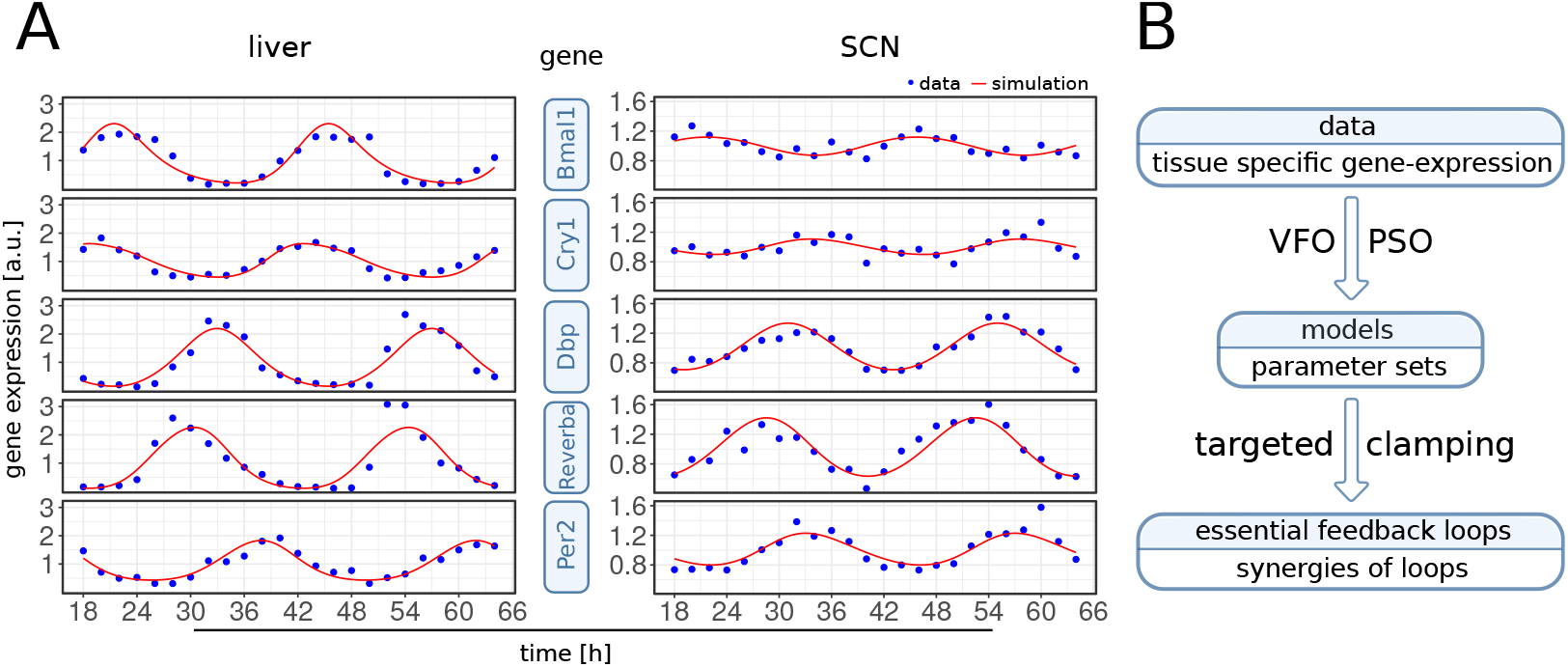
(A) Example time series for data and simulation. One fit in liver (left) and one fit in the SCN (right) are shown. Expression levels are normalized to the mean values. The liver fit has a score of 0.01 and involves the *Bmal1-Rev-erb-α*, repressilator and *Cry1* loops, while the SCN fit scores 3.36 and involves the *Bmal1-Rev-erb-α, Per2* and *Cry1* loops. Note the smaller amplitudes and the early *Cry1* phase in the SCN. (B) Workflow of the analysis. Multiple optimized parameter sets are obtained from each tissue-specific data set. Then essential loops are identified in the respective models.

### 2.2 Vector field optimization improves model fitting

To investigate whether our reduced gene regulatory network can reproduce tissue-specific data [3] we developed a pipeline for global parameter optimization. We applied the pipeline multiple times to each tissue-specific expression profile, allowing us to compare optimized model parameters between tissues. Table 1 lists the number of optimization runs for 10 analyzed tissues. Four tissues (liver, SCN, adrenal gland and kidney) are discussed in more detail in the following sections, while results for others can be found in Supplements S2 and S5.

**Table 1:** Number of optimization runs per tissue and average score. Four tissues are mainly discussed in the main text and others are described in supplements.

|  | adrenal gland | kidney | liver | heart | skeletal muscle | lung | brown adipose | white adipose | SCN | cerebellum |
| --- | --- | --- | --- | --- | --- | --- | --- | --- | --- | --- |
| number of runs | 100 | 93 | 57 | 57 | 58 | 31 | 62 | 45 | 153 | 58 |
| runs with score < 10 | 66 | 59 | 52 | 44 | 35 | 21 | 36 | 22 | 46 | 39 |
| mean score | 3.84 | 4.48 | 1.58 | 3.74 | 5.17 | 3.04 | 4.00 | 3.24 | 7.21 | 3.99 |
| Figure | 2,5,6,7,8 | 2,5,6,7,8 | 2-8 | 8, S2,S5 | 8, S2,S5 | 8, S2,S5 | 8, S2,S5 | 8, S2,S5 | 2-8 | S2,S5 |

The agreement of model simulations and experimental mRNA time courses [3] was measured by a scoring function. In this function period, relative phases and fold changes of measured gene transcripts are taken into account. The complete scoring function is given in Supplement S3.

Model parameters are chosen by global optimization, such that the score obtained by our scoring function is minimal. The optimization method approaches a local minimum in a high dimensional parameter space and thus final scores of each run depend on the starting conditions. We only used model fits with scores below a chosen threshold of 10 for further analyses (see Supplement S3). Interestingly, the fractions of optimization runs with scores below 10 vary across tissues. While the largest number of successful runs is found for liver data (about 90 %), for kidney and adrenal gland about 2/3 and for SCN only 1/3 of the runs yield good scores below the chosen threshold.

Allowed ranges for parameters were defined to restrict the search space. While delays and degradation rates are optimized within biologically plausible ranges around experimentally measured values, for activation and inhibition strengths no such measurements are available. Therefore, we define ranges based on oscillation mean levels and corresponding to the working points of regulations, that is ranges in parameter space in which regulation strengths vary most.

Global optimization is performed with Particle Swarm Optimization (PSO) [38]. A number of particles—each representing one parameter combination—is initialized randomly using latin hypercube sampling [39] and moved around in the parameter space with velocities changing according to both their individual and their neighbors known best location. The movements are conducted for a number of iterations while velocities decrease and particles converge to an optimum.

We improve global optimization by identifying good starting conditions. To this end, we devise a strategy which we here call “vector field optimiza-tion” (VFO). Our algorithm makes use of available time courses for each variable and their mathematical description in terms of differential equations. From the data, we can approximate the time derivatives together with the right hand sides of our model equations (Fig. 4A, Supplement S4). By minimizing the differences we obtain initial values of model parameters. This step does not require simulation, but already yields parameter combinations that account for much of the differences between time courses. For example, the known antiphase oscillations of *Bmal1* and *Rev-erb-α* can be generated with a *Bmal1*-delay of about 6 h. Even though the overall search space of this parameter is the interval from 0 to 6 hours, VFO leads to initial delay values close to 6 h (see Supplement 4 for details).

VFO is performed using a bounded gradient method to ensure that solutions lie within the parameter limits. Starting points for the gradient method are chosen randomly. We tested whether VFO improves the scores of model fits. Indeed, we are able to find significantly more good fits for liver and SCN then with PSO alone (Figure 4B). Notably, in the SCN it was difficult to reach scores below 10 without previous application of VFO.

**Figure 4:**
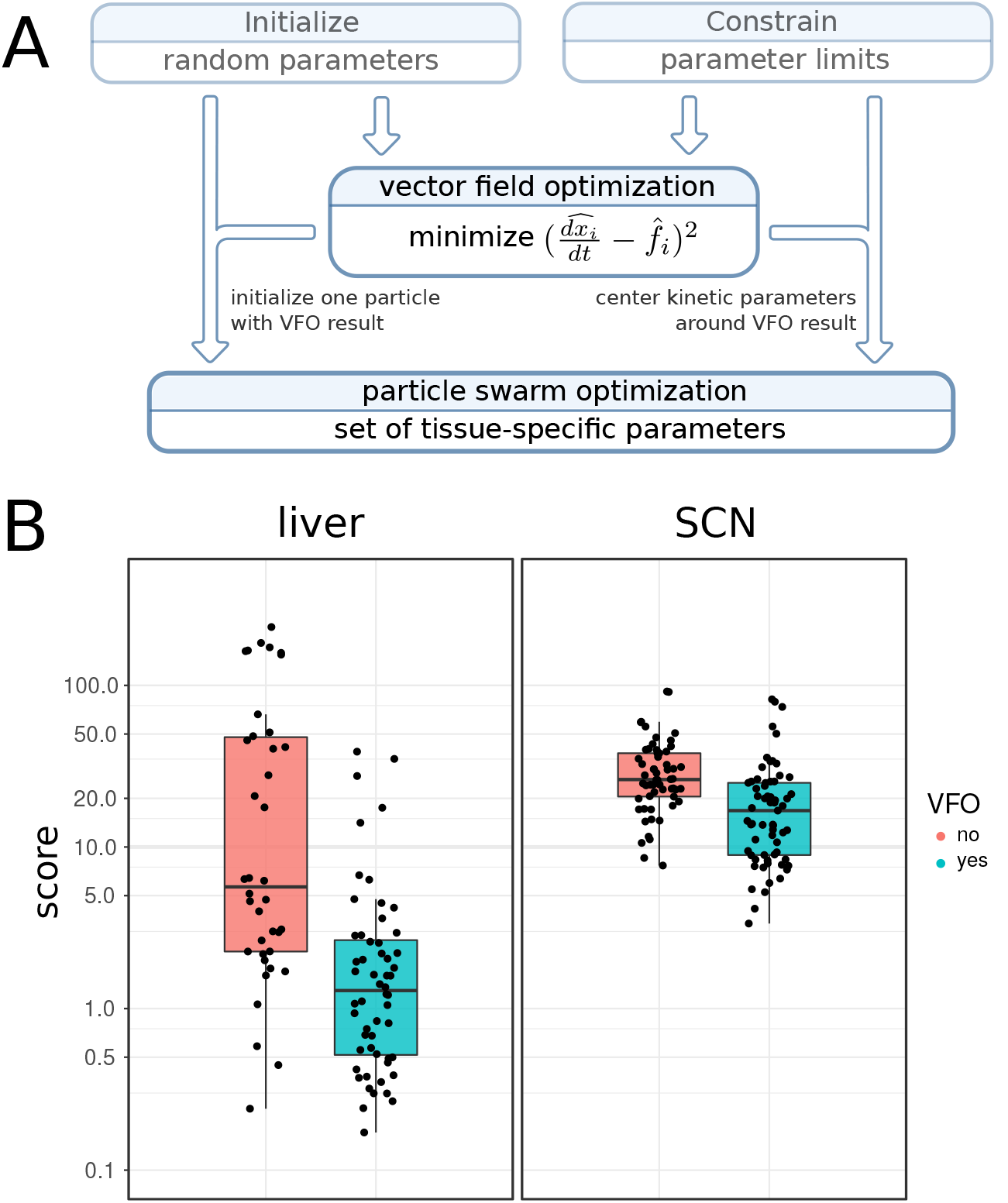
Improved fitting with vector field optimization (VFO). (A) Flow diagram showing how VFO is integrated into the fitting procedure. The resulting parameter set is used to initialize one particle and to pre-emphasize kinetic parameters. (B) Score for fits to circadian transcription data from mouse liver and SCN with and without vector field optimization. Each point represents a fitted model. VFO leads to significantly lower score values (Wilcoxon rank sum test, p-value liver: 4.29· 10^−7^, p-value SCN: 4.819· 10^−6^).

### 2.3 Clamping reveals essential loops

Using global optimization we found for all 10 tissue-specific expression profiles [3] parameter sets that reproduce the data within experimental uncertainties (Supplement S2 and S3). There were not just single optima of global optimization, but for all investigated tissues multiple parameter configurations fitted the data.

To determine essential feedback loops for each model fit, we use our clamping protocol published in 2016 [13]. Clamping of genes is done by setting the expression level of genes to their mean value (constant) and corresponds to constitutive expression experiments in the wet lab [36, 40–42]. It allows comparison of the effect of rhythmic versus basal regulation.

In addition to gene clamping we also clamp specific regulations via gene products. In silico this is done by setting the corresponding terms in the differential equations of the model constant. Regulations/terms are shown as network links in Figure 1. We can examine the relevance of feedback loops associated with such links by clamping regulations systematically.

To reduce computational effort we use a targeted clamping strategy, testing specifically which feedback loops are essential. We regard a negative feedback loop as essential for oscillations if clamping of each link that is part of the loop disrupts of rhythmicity (only one link at a time is clamped). Details are provided in Supplement S5.

In addition we test the synergy of loops by clamping combinations of regulations. We are able to distinguish two different modes of synergistic function: (A) two loops work independent of each other and mutually compensate for perturbations and (B) two dependent loops share the required feedback for oscillations, such that rhythms only occur when both loops are active.

For example, in liver we found many model fits in which the *Bmal1-Rev-erb-α* and repressilator loops function synergistically. Both loops constitute a negative feedback from *Rev-erb-α* onto itself and are timed accordingly, such that they mutually support rhythmicity.

We apply this clamping analysis to every generated model fit (420 in total, see Table 1) to identify feedback loops responsible for rhythmicity. Most commonly found loops are marked in Figure 1.

### 2.4 Loop- and parameter composition reflects variation of clock gene expression

For every tissue-specific data set we have obtained multiple model fits. After assigning essential loops to every model fit, we are able to compare variations in loop composition between different tissues.

Therefore, we define four core loops that were identified by our analysis and discussed in the literature. Figure 5 shows how the composition of these four loops varies between tissues. Interestingly, there are marked differences.

**Figure 5:**
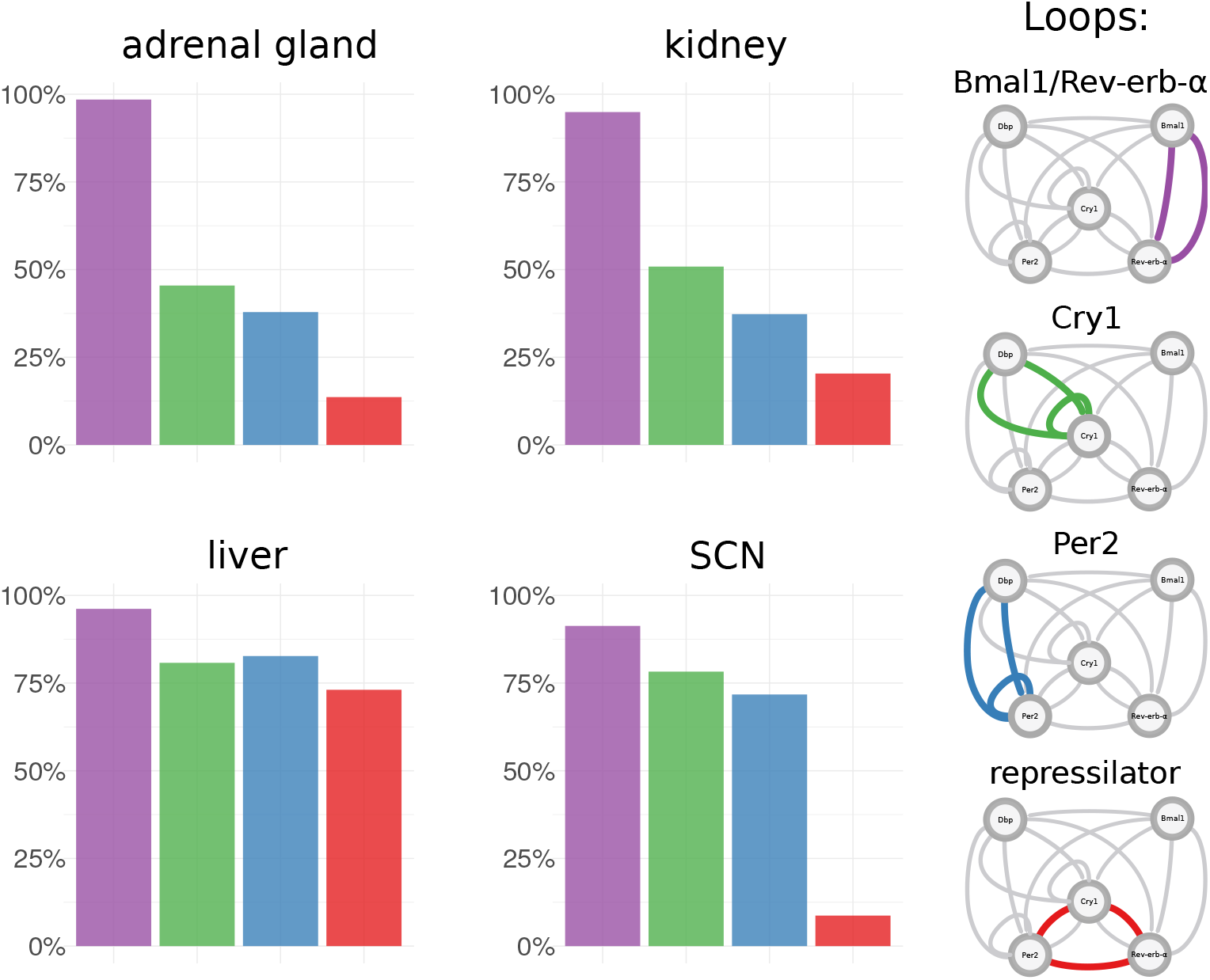
Proportions of essential feedback loops across tissues. For each tissue-specific transcriptome data set multiple model fits were generated. In each model fit essential feedback loops were then identified using clamping analysis. Frequencies are shown for a set of four core loops that were most prominent in the analysis and are discussed in the literature.

In liver *Bmal1-Rev-erb-α, Per2, Cry1* loops and repressilators all occur with comparable frequencies and thus appear to fit the data equally well. In contrast, in the SCN repressilators only occur in a few cases. This is also consistent with the early peak time of *Cry1* mentioned earlier, since the re-pressilator mechanism is based on distinct inhibitions at different phases [13]. Fits to adrenal gland and kidney data have similar proportions of essential loops and in contrast to liver they have more *Bmal1-Rev-erb-α* loops and less repressilators. The profiles of additional tissues are shown in Supplement 5.

To find out whether tissue differences are reflected in the model parameters, we examine their distributions in the 34-dimensional parameter space. To this end, we perform dimensionality reduction by principal component analysis (PCA) and visualize tissue-differences using Linear Discriminant Analysis (LDA) [43].

Figure 6A illustrates that the parameters sets fitted to SCN are clearly different from those fitted to other tissues. The differences can be assigned to selected parameters as indicated by the red arrows.

**Figure 6:**
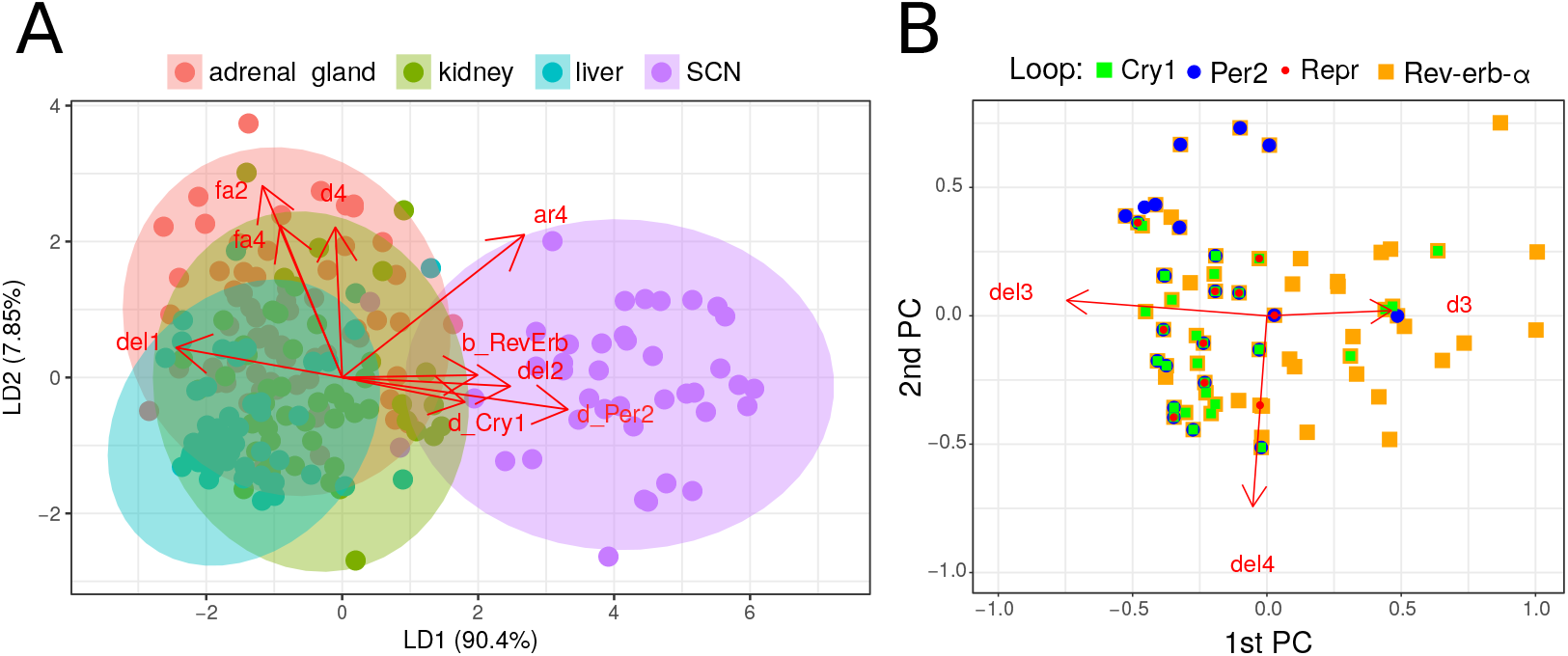
Tissue-specific models separated in parameter space. (A) Linear Discriminant Analysis. Fits (points) are projected to a plane while trying to maximize the variance between tissues. The projected parameter vectors are visualized as arrows in this plane. The delays of *Bmal1* (del1), *Rev-erb-α* (del2) and degradation rates of *Per2* (d_Per2), *Cry1* (d_Cry1) and *Rev-erb-α* (d_RevErb) as well as the *Cry1* inhibition by *Rev-erb-α* (ar4) differ most between SCN fits and fits of other tissues. (B) Loops in parameter space. Shown are the first two principal components and points corresponding to parameter sets for adrenal gland. Directions of the parameter axes are given as red arrows. Relations between parameter values and loops are visible, for example essential *Cry1* loops (green) occur, when *Cry1* delays (del4) are large.

In Figure 6B we project the model parameters to the first two principal components and color the points according to the essential loops. It turns out that *Per2* loops (blue), *Cry1* loops (green) and *Bmal1/Rev-erb-α* loops (orange) are associated with distinct parameter sets. We observe, for example, an association of *Cry1* loops with high *Cry1* delay.

The differences in loop distributions (Figure 5) and parameter constellations (Figure 6) suggest that differences between expression profiles (see Figure 2) imply tissue-specific mechanisms to generate self-sustained oscillations. For example, in brain tissues small amplitudes and early *Cry1* phases promote self-inhibitions of *Per2* and *Cry1* whereas a large *Rev-erb-α* amplitude in liver leads to many solutions with *Bmal1/Rev-erb-α* loops and repressilators.

### 2.5 Synergies of feedback loops

So far, we discussed tissue-specific frequencies of single loops. Interestingly, most parameter sets cannot be assigned to unique loops but to combinations of different essential feedback loops. Now we use a targeted clamping strategy (Supplement S5) to explore possible synergies of feedback loops.

Our clamping strategy allows us to find loops that are necessary (or essential) for rhythm generation. If we clamp regulations that are part of these loops, rhythms vanish. Further by clamping many regulations at the same time, we can also identify sets of loops that are sufficient for oscillation generation. In simulations rhythms persist, if these loops are active while all others are clamped. We term such synergistic sets of loops “rhythm generating oscillators”.

Analyzing 420 parameter sets we find more than 70% that exhibit synergies of different feedback loops. Figure 7A illustrates that most models constitute combinations of loops. Venn diagrams in Figure 7B show that in liver *Bmal1-Rev-erb-α* loops together with repressilators form the largest group of oscillators whereas in the SCN *Bmal1-Rev-erb-α* loops are typically associated with *Per2* and *Cry1* loops.

**Figure 7:**
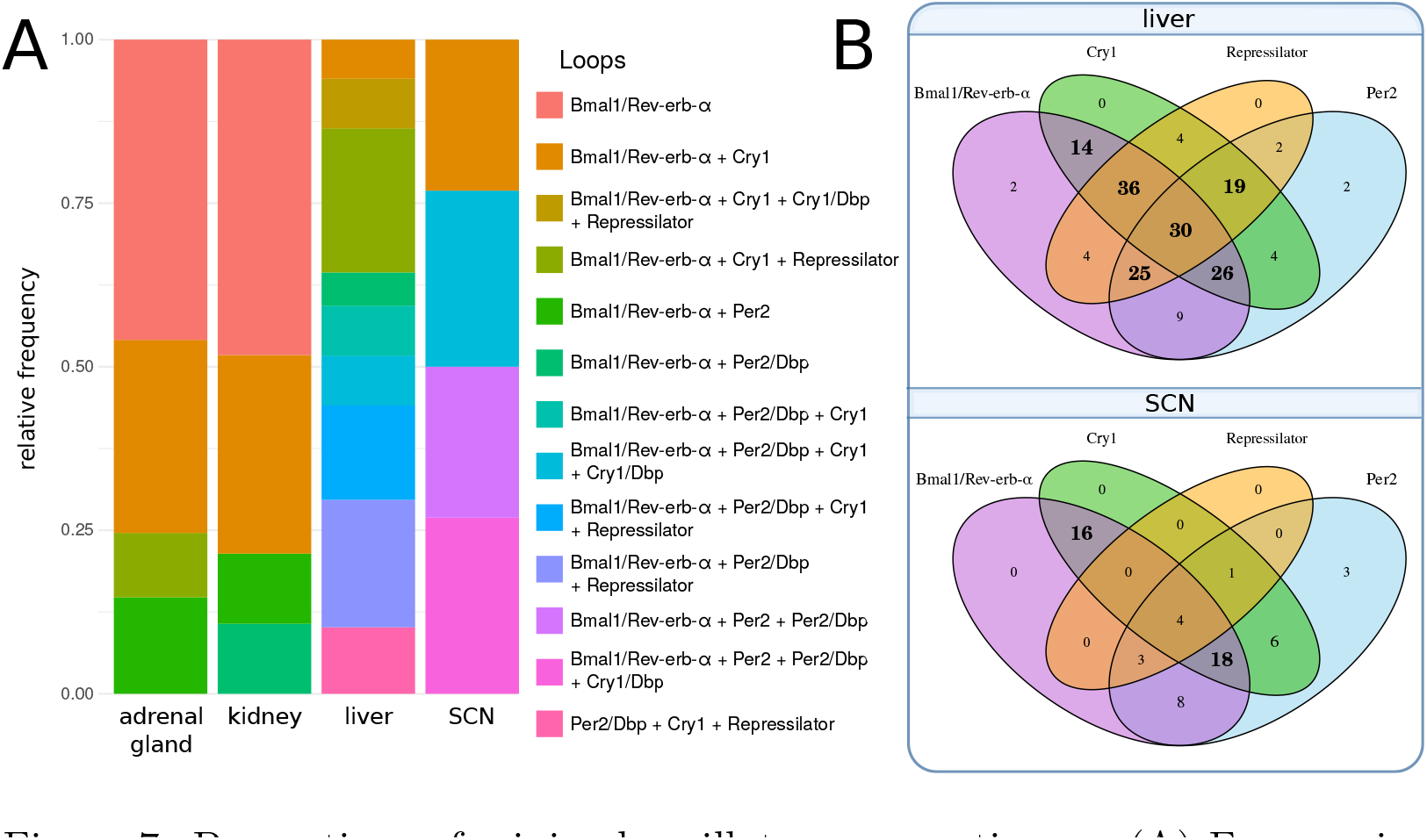
Proportions of minimal oscillators across tissues. (A) Frequencies of oscillators in models of different tissues. An oscillator comprises one or more loops (connected with a “+” in the legend). *Per2, Cry1* loops and their extensions via *Dbp* (see Figure 1) are counted separately here. (B) Venn diagrams of oscillator composition for liver and SCN. Bold numbers highlight the most frequent subsets. In liver most oscillators comprise many loops including *Bmal1/Rev-erb-α* loop, repressilator, *Per2* and *Cry1 loops*. In the SCN most oscillators are a combination of *Bmal1/Rev-erb-α* with *Cry1* and *Per2*.

Interestingly, the synergy of multiple loops leads typically to low scores. There is a significant difference in the number of loops between parameter sets above and below the median score (Wilcoxon rank-sum test, p-value < 0.0001) and most model fits involving all four loops lead to excellent scores below 2.5.

Moreover, repressilators exhibit quite good scores in particular for liver, kidney, and adrenal gland. For these tissues fits with repressilator have an average score of 1.24, while fits without repressilator have a mean score of 4.41. This is consistent with our finding that better scores involve more loops. The repressilator motif connects inhibitions of *Per2, Cry1* and *Rev-erb-α* and links that way loops synergistically.

### 3 Discussion

Circadian rhythms in mammals are generated by a cell-autonomous gene regulatory network [44]. About 20 regulators drive core clock genes via E-boxes, D-boxes and ROR-elements [10]. Based on clustered gene expression phases (compare Fig. 2) we reduced the system to a network of 5 genes connected by 7 positive and 10 negative regulations (see Fig. 1). This reduced model still contains multiple positive and negative feedback loops.

Our aim was to identify the most essential feedback loops and to quantify tissue differences. Our network model was fitted to comprehensive expression profiles of 10 mammalian tissues [3]. Furthermore, we use proteomics data [22], ChIP-seq data [11, 23], and decay-rate data [25] to constrain the ranges of unknown parameters. Since quantitative data on protein dynamics are sparse, we simplified the model by using explicit delays between transcription and regulation.

We optimized parameters by a combination of a novel approach termed Vector Field Optimization (VFO) and Particle Swarm Optimization (PSO) [38]. After combining these global optimization techniques our simulations could reproduce the data within experimental uncertainties.

To our surprise, we found for every studied tissue multiple excellent fits with quite different parameter constellations. In order to extract the responsible feedback loops we performed a systematic clamping analysis. Individual regulatory terms (edges in the network) were systematically clamped to constant values. These clamping methods revealed the essential feedback loops in each of the networks derived from tissue-specific expression-profiles.

We found an astonishing diversity of essential feedback loops in models that were able to reproduce the experimental data. Among the essential loop structures we found *Per* and *Cry* self-inhibitions. These loops have been considered as the primary negative feedacks since the double knockouts of *Cry* genes [45] and the triple knockout of *Per2* genes [46] were arrhythmic. Later additional feedback loops via nuclear receptors have been found [47]. As predicted by modeling [17] and confirmed by *Rev-erb* double knockouts [18] the *Bmal1-Rev-erb* loops constitute another possible rhythm generator. Indeed, in all tissues we detected parameter constellations that employ this negative feedback loop.

Recently, the repressilator, a chain of serial inhibitions, was suggested as possible mechanism to generate oscillations in liver and adrenal gland [13]. This loop structure is associated with dual modes of E-box inhibitions [30] based late *Cry1* expression [35] and late CRY1 binding to E-boxes [11]. Since the expression phase of *Cry1* is tissue-dependent it is plausible that also the detection of repressilators might differ between different organs.

Indeed, repressilators are less frequently detected as essential in models based on brain data-derived parameter sets as shown in Fig. 5. In general, the model parameters in the SCN are clearly different from the parameters in peripheral tissues (see Figure 6). Thus, modeling can point to different design principles in specific organs. The large amplitudes and late *Cry1* phases in tissues such as liver suggest that the repressilator is a relevant mechanism in these tissues whereas small amplitudes and early *Cry1* phase in brain tissues favor *Cry* and *Per* self-inhibitions.

Major differences between tissues have been reported also regarding amplitudes and phases of clock-controlled genes [3-6]. Such differences are presumably induced by tissue-specific transcription factors [34, 48, 49]. Moreover, different organs receive different metabolic and neuroendocrine inputs leading to quite different rhythmic transcriptomes. These systemic tissue differences can also modify the core clock dynamics. In particular nuclear receptor rhythms differ drastically between organs [50] and can induce tissue-specificities of the core clocks [51, 52].

Interestingly, the best scoring models involve several essential feedback loops. This observation indicates that the synergy of different feedback mechanisms improves oscillator quality. Furthermore, co-existing loops imply redundancy and thus the core clock is buffered with respect to non-optimal gene expression, hormonal rhythms, seasonal variations and environmental fluctuations.

Co-occuring feedback regulations might also explain reports where different circadian outputs displayed slightly different periods. For example, in SCN slices different reporter signals indicated distinct periods [53, 54] and also in crickets two independent negative feedback loops were reported [55]. In some of our high-scoring networks we find indeed two independent frequencies leading to slight modulations of the circadian waveforms (see Supplement S6).

Tissue-specific core clock mechanisms are presumably related to functional differences of SCN and peripheral organs. *Per* gene regulations are particularly important in the SCN since light inputs and coupling via VIP induce *Per* genes via CREB [16, 56]. Relatively small core clock amplitudes in the SCN allow efficient entrainment and synchronization [57, 58]. Moreover, small amplitudes might facilitate adaptation to long and short photoperiods by varying coupling mechanisms [59, 60]. The dominant role of *Per* and *Cry* self-inhibitions is also reflected by arrhythmic activities of *Per* and *Cry* double knockouts [45, 46].

Peripheral organs such as liver, kidney, and adrenal gland govern the daily hormonal and metabolic rhythms. Consequently, large amplitudes and pronounced rhythms of nuclear receptors are observed [50, 61]. Interestingly, we find that feedback loops involving ROR-elements are more prominent in these tissues (compare Fig. 8).

In summary, our study suggests that there is not necessarily a single dominant feedback loop in the mammalian core clock. Instead, multiple mechanisms including *Per/Cry* self-inhibitions, *Bmal1-Rev-erb* loops, and repressilators are capable to generate circadian rhythms. The co-existence of feedback loops provides redundancy and can thus enhance robustness and flexibility of the intertwined circadian regulatory system.

## Acknowledgements

We are grateful for discussions with Dr. Christoph Schmal, Dr. Bharath Ananthasubramaniam, Dr. Anja Korenčič and Dr. Matthias König. This work was supportet by grants from Deutsche Forschungsgemeinschaft (HE2168/11-1, TRR/SFB 186 A16 and A17), BMBF (01GQ1503) and GRK CSB 1772/2.

## S1 Model equations

### Underlying mathematical model

For fitting experimental data we start with a network model that was introduced by Korenčič et al. in 2014 [1]. It uses only 5 variables representing groups of genes with distinct expression peaks. Genes with the same peak time and binding mechanism of their protein products are modeled by the same variable. For example, all early E-box inhibitors, such as *Per1, Per2, Per3* and *Cry2* are represented by a single variable called *Per2*. The late E-box inhibitor *Cry1*, however, is modeled as a second variable.

Further, the model consists of Delay-Differential Equations (DDE). Regulations affects other genes only after certain time delays, which are parameters in the model. A time delay could correspond to time consuming processes such as nuclear translocation or complex formation and is important for oscillations with a circadian period. Models using Ordinary Differential Equations (ODE) require additional intermediate steps to obtain the right temporal scales. Thus, use of DDE also allows to represent the core clock network with a small number of variables and parameters.

Taken together, the model constitutes a condensed representation of the core clock network. It is not targeted at capturing all mechanistic details, but rather presents a regression that exploits timing of gene expression to understand self-sustained circadian oscillations.

### Structure of Equations

The full set of equations is shown in Example S1-1. Note the modular structure of activatory and inhibitory terms, marked with blue and red labels respectively. Note, that activator terms contain two free parameters (maximum fold induction and activation threshold), whereas inhibitory terms have just a single free parameter—the inhibition threshold. The interactions of the model were assembled in a comprehensive survey of ChIP-Seq experiments and motif mapping. The number of binding sites of a gene is reflected in the power of the corresponding term.

---

**Example S1-1.**
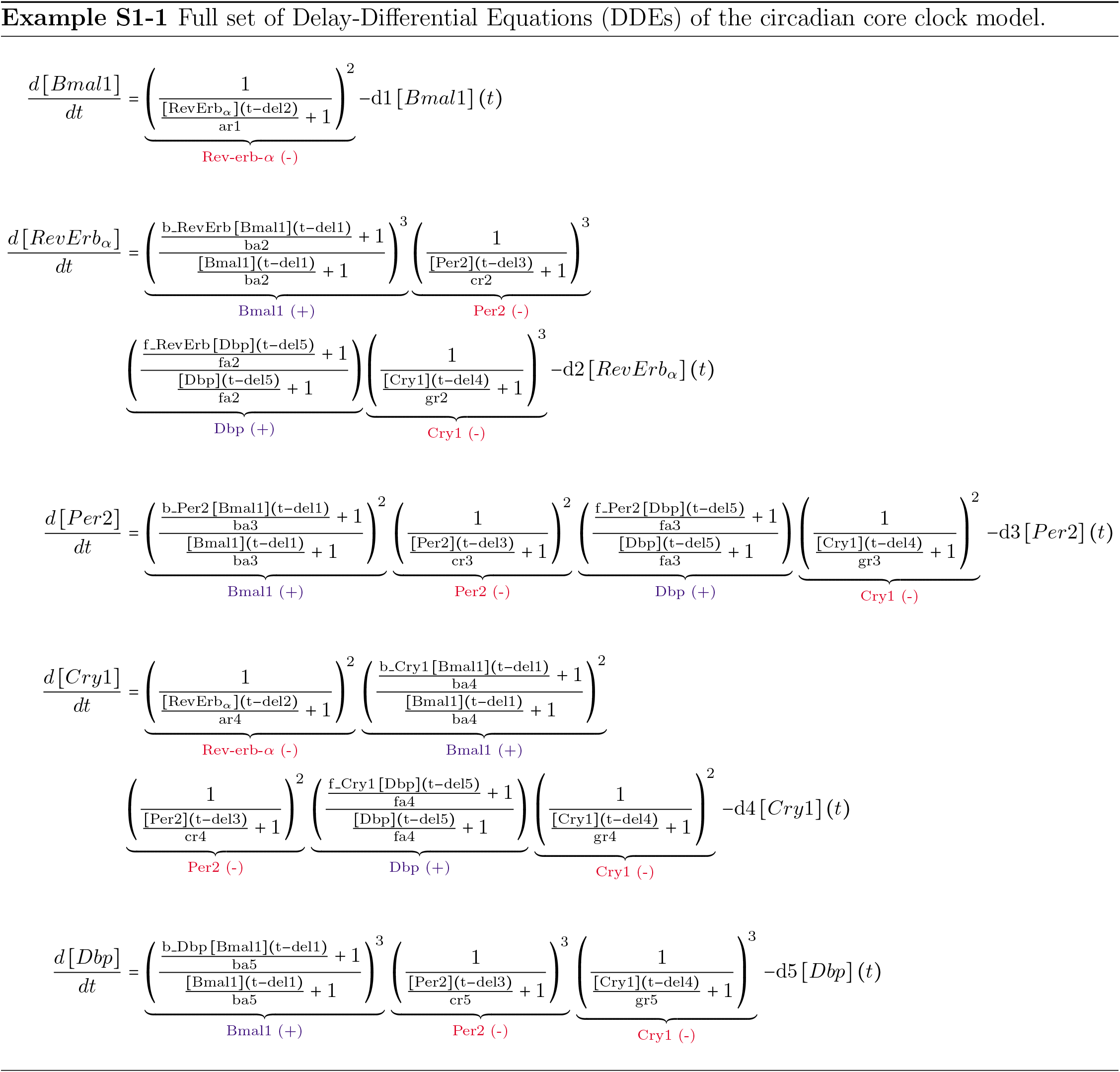
Full set of Delay-Differential Equations (DDEs) of the circadian core clock model.

## S2 Parameter sets

Here we list sample parameter sets for each tissue with their score and minimal oscillators. The score was given according to the scoring function in Supplement S3. “Minimal oscillator” refers to the finding, that the listed loops can synergistically generate oscillations, if they are not clamped. Minimal means that at least these loops need to be active. The models shown here are taken from the top scoring fits of each tissue.

### adrenal gland

**score:** 0.091
**oscillator:***Bmal1/Rev-erb-α* + Repressilator

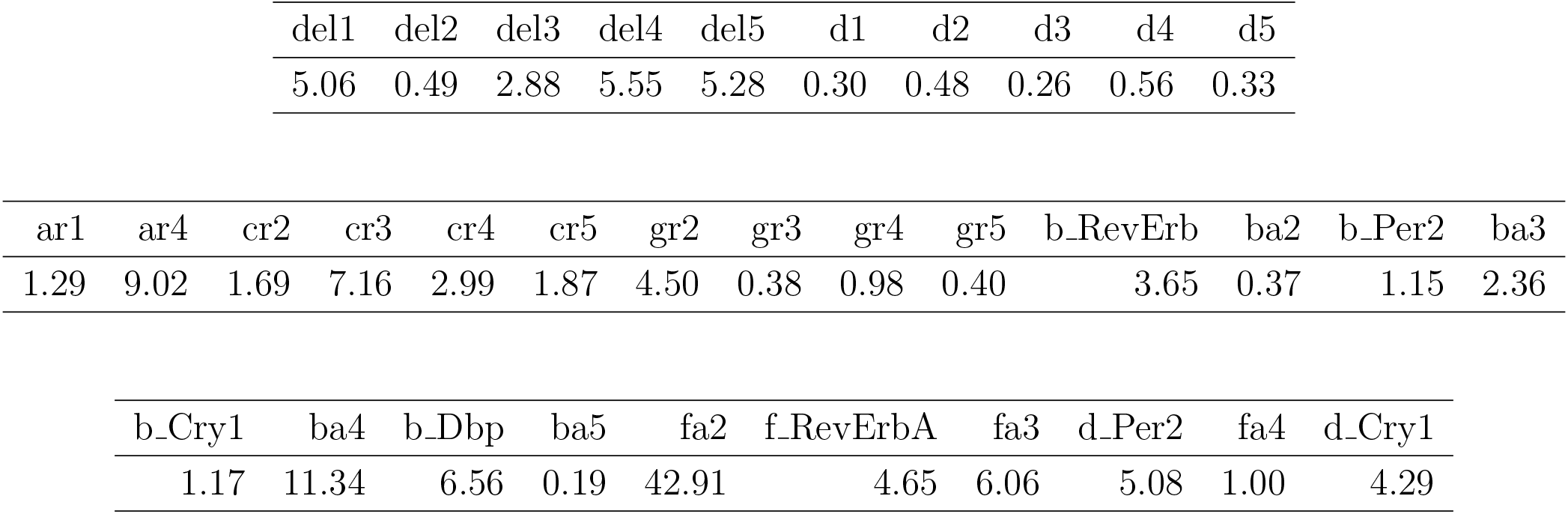

### kidney

**score:** 0.090
**oscillator 1:** Bmal1/Reverba + Per2 + Per2/Dbp + Repressilator
**oscillator 2:** Bmal1/Reverba + Per2/Dbp + Cry1 + Repressilator
**oscillator 3:** Bmal1/Reverba + Per2/Dbp + Cry1/Dbp + Repressilator

| del1 | del2 | del3 | del4 | del5 | d1 | d2 | d3 | d4 | d5 |
| --- | --- | --- | --- | --- | --- | --- | --- | --- | --- |
| 5.65 | 0.88 | 5.34 | 4.81 | 4.50 | 0.50 | 0.73 | 0.41 | 0.27 | 0.43 |

| ar1 | ar4 | cr2 | cr3 | cr4 | cr5 | gr2 | gr3 | gr4 | gr5 | b_RevErb | ba2 | b_Per2 | ba3 |
| --- | --- | --- | --- | --- | --- | --- | --- | --- | --- | --- | --- | --- | --- |
| 1.51 | 2.99 | 1.92 | 4.82 | 1.95 | 0.71 | 4.04 | 0.23 | 0.19 | 0.42 | 2.69 | 0.34 | 1.50 | 9.62 |

| b_Cry1 | ba4 | b_Dbp | ba5 | fa2 | f_RevErbA | fa3 | d_Per2 | fa4 | d_Cry1 |
| --- | --- | --- | --- | --- | --- | --- | --- | --- | --- |
| 1.23 | 0.82 | 3.50 | 0.04 | 12.02 | 2.09 | 4.22 | 5.87 | 1.55 | 4.25 |

### liver

**score:** 0.014
**oscillator 1:** Bmal1/Reverba + Per2/Dbp + Repressilator
**oscillator 2:** Bmal1/Reverba + Cry1 + Repressilator

| del1 | del2 | del3 | del4 | del5 | d1 | d2 | d3 | d4 | d5 |
| --- | --- | --- | --- | --- | --- | --- | --- | --- | --- |
| 5.83 | 0.74 | 2.74 | 3.35 | 5.29 | 0.29 | 0.77 | 0.42 | 0.23 | 0.42 |

| ar1 | ar4 | cr2 | cr3 | cr4 | cr5 | gr2 | gr3 | gr4 | gr5 | b_RevErb | ba2 | b_Per2 | ba3 |
| --- | --- | --- | --- | --- | --- | --- | --- | --- | --- | --- | --- | --- | --- |
| 1.83 | 4.40 | 0.84 | 14.82 | 12.16 | 3.16 | 8.03 | 0.87 | 1.83 | 0.67 | 4.06 | 0.12 | 1.81 | 3.45 |

| b_Cry1 | ba4 | b_Dbp | ba5 | fa2 | f_RevErbA | fa3 | d_Per2 | fa4 | d_Cry1 |
| --- | --- | --- | --- | --- | --- | --- | --- | --- | --- |
| 1.66 | 0.52 | 4.40 | 0.90 | 12.68 | 2.61 | 1.74 | 4.03 | 4.15 | 5.59 |

### SCN

**score:** 3.49
**oscillator:** Per2/Dbp + Cry1 + Cry1/Dbp

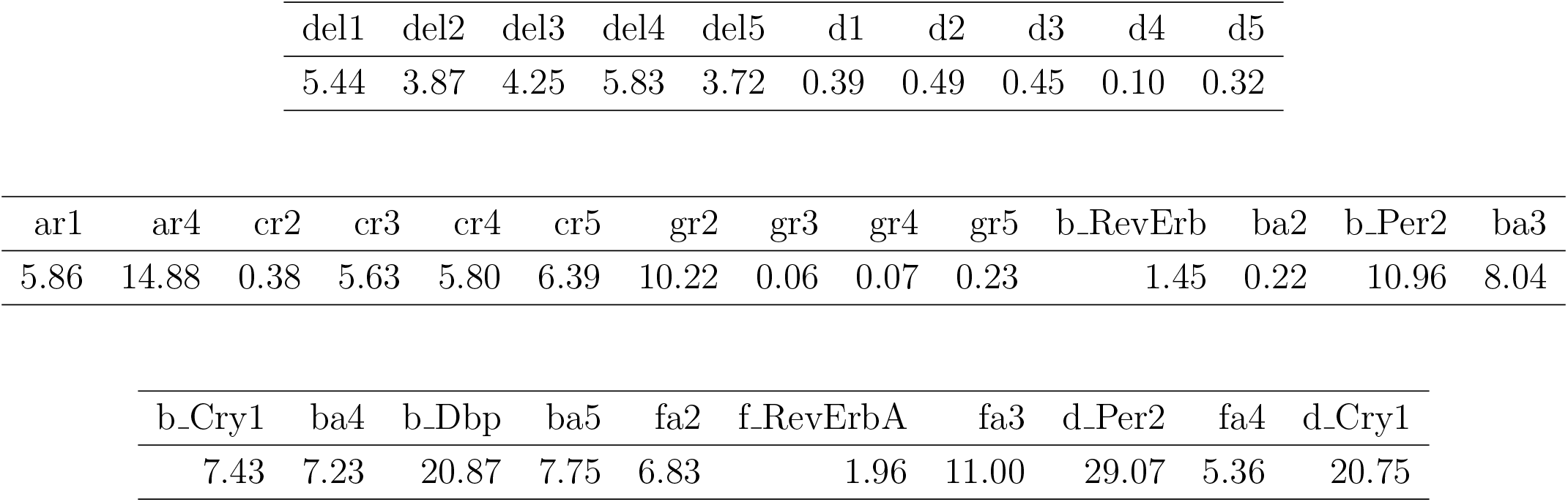

### heart

**score:** 0.19
**oscillator:** Bmal1/Reverba

| del1 | del2 | del3 | del4 | del5 | d1 | d2 | d3 | d4 | d5 |
| --- | --- | --- | --- | --- | --- | --- | --- | --- | --- |
| 5.62 | 1.47 | 3.87 | 2.37 | 4.46 | 0.36 | 0.46 | 0.95 | 0.22 | 0.92 |

| ar1 | ar4 | cr2 | cr3 | cr4 | cr5 | gr2 | gr3 | gr4 | gr5 | b_RevErb | ba2 | b_Per2 | ba3 |
| --- | --- | --- | --- | --- | --- | --- | --- | --- | --- | --- | --- | --- | --- |
| 3.52 | 20.43 | 9.97 | 5.68 | 11.03 | 2.56 | 4.09 | 0.11 | 0.18 | 0.05 | 5.71 | 1.00 | 4.39 | 7.50 |

| b_Cry1 | ba4 | b_Dbp | ba5 | fa2 | f_RevErbA | fa3 | d_Per2 | fa4 | d_Cry1 |
| --- | --- | --- | --- | --- | --- | --- | --- | --- | --- |
| 3.69 | 12.35 | 10.62 | 0.07 | 21.22 | 2.85 | 15.87 | 16.60 | 4.75 | 5.31 |

### skeletal muscle

**score:** l.l
**oscillator 1:** Bmal1/Reverba + Per2
**oscillator 2:** Bmal1/Reverba + Per2 + Per2/Dbp
**oscillator 3:** Bmal1/Reverba + Per2 + Cry1
**oscillator 4:** Bmal1/Reverba + Per2/Dbp + Cry1

| del1 | del2 | del3 | del4 | del5 | d1 | d2 | d3 | d4 | d5 |
| --- | --- | --- | --- | --- | --- | --- | --- | --- | --- |
| 4.08 | 4.58 | 5.98 | 4.76 | 3.37 | 0.72 | 0.94 | 0.35 | 0.37 | 0.16 |

| ar1 | ar4 | cr2 | cr3 | cr4 | cr5 | gr2 | gr3 | gr4 | gr5 | b_RevErb | ba2 | b_Per2 | ba3 |
| --- | --- | --- | --- | --- | --- | --- | --- | --- | --- | --- | --- | --- | --- |
| 5.54 | 18.49 | 12.63 | 1.15 | 6.96 | 0.19 | 5.32 | 0.94 | 2.89 | 11.30 | 13.03 | 4.74 | 1.00 | 14.65 |

| b_Cry1 | ba4 | b_Dbp | ba5 | fa2 | f_RevErbA | fa3 | d_Per2 | fa4 | d_Cry1 |
| --- | --- | --- | --- | --- | --- | --- | --- | --- | --- |
| 3.60 | 9.70 | 11.95 | 0.18 | 6.88 | 4.22 | 30.74 | 21.44 | 32.56 | 13.75 |

### brown adipose

**score:** 0.276
**oscillator:** Bmal1/Reverba

| del1 | del2 | del3 | del4 | del5 | d1 | d2 | d3 | d4 | d5 |
| --- | --- | --- | --- | --- | --- | --- | --- | --- | --- |
| 5.60 | 0.52 | 2.92 | 4.27 | 4.77 | 0.23 | 0.84 | 0.71 | 0.28 | 0.32 |

| ar1 | ar4 | cr2 | cr3 | cr4 | cr5 | gr2 | gr3 | gr4 | gr5 | b_RevErb | ba2 | b_Per2 | ba3 |
| --- | --- | --- | --- | --- | --- | --- | --- | --- | --- | --- | --- | --- | --- |
| 2.48 | 87.53 | 3.71 | 2.79 | 0.19 | 1.67 | 9.08 | 0.15 | 2.96 | 0.44 | 10.62 | 0.88 | 1.04 | 6.23 |

| b_Cry1 | ba4 | b_Dbp | ba5 | fa2 | f_RevErbA | fa3 | d_Per2 | fa4 | d_Cry1 |
| --- | --- | --- | --- | --- | --- | --- | --- | --- | --- |
| 1.38 | 7.26 | 22.54 | 0.14 | 22.34 | 3.75 | 10.43 | 4.13 | 10.39 | 4.25 |

### white adipose

**score:** 0.171
**oscillator1:** Bmal1/Reverba + Per2/Dbp + Repressilator
**oscillator2:** Bmal1/Reverba + Cry1 + Repressilator

| del1 | del2 | del3 | del4 | del5 | d1 | d2 | d3 | d4 | d5 |
| --- | --- | --- | --- | --- | --- | --- | --- | --- | --- |
| 5.88 | 2.05 | 5.55 | 3.36 | 4.67 | 0.85 | 0.82 | 0.39 | 0.41 | 0.29 |

| ar1 | ar4 | cr2 | cr3 | cr4 | cr5 | gr2 | gr3 | gr4 | gr5 | b_RevErb | ba2 | b_Per2 | ba3 |
| --- | --- | --- | --- | --- | --- | --- | --- | --- | --- | --- | --- | --- | --- |
| 2.96 | 7.38 | 1.70 | 4.73 | 14.88 | 0.25 | 3.07 | 0.24 | 0.31 | 1.89 | 3.46 | 0.45 | 2.10 | 6.46 |

| b_Cry1 | ba4 | b_Dbp | ba5 | fa2 | f_RevErbA | fa3 | d_Per2 | fa4 | d_Cry1 |
| --- | --- | --- | --- | --- | --- | --- | --- | --- | --- |
| 1.04 | 5.69 | 8.07 | 0.06 | 3.43 | 1.51 | 6.83 | 6.44 | 3.31 | 2.79 |

### lung

**score:** 0.34
**oscillator:** Bmal1/Reverba

| del1 | del2 | del3 | del4 | del5 | d1 | d2 | d3 | d4 | d5 |
| --- | --- | --- | --- | --- | --- | --- | --- | --- | --- |
| 5.50 | 0.59 | 3.01 | 2.88 | 5.32 | 0.18 | 0.78 | 0.45 | 0.23 | 0.34 |

| ar1 | ar4 | cr2 | cr3 | cr4 | cr5 | gr2 | gr3 | gr4 | gr5 | b_RevErb | ba2 | b_Per2 | ba3 |
| --- | --- | --- | --- | --- | --- | --- | --- | --- | --- | --- | --- | --- | --- |
| 1.11 | 29.82 | 1.60 | 10.51 | 9.90 | 9.73 | 7.46 | 0.24 | 0.22 | 0.40 | 7.21 | 0.54 | 1.84 | 7.92 |

| b_Cry1 | ba4 | b_Dbp | ba5 | fa2 | f_RevErbA | fa3 | d_Per2 | fa4 | d_Cry1 |
| --- | --- | --- | --- | --- | --- | --- | --- | --- | --- |
| 2.02 | 9.49 | 12.53 | 0.59 | 11.62 | 3.01 | 4.32 | 7.17 | 1.23 | 6.63 |

### cerebellum

**score:** 0.545
**oscillator:** Bmal1/Reverba + Cry1

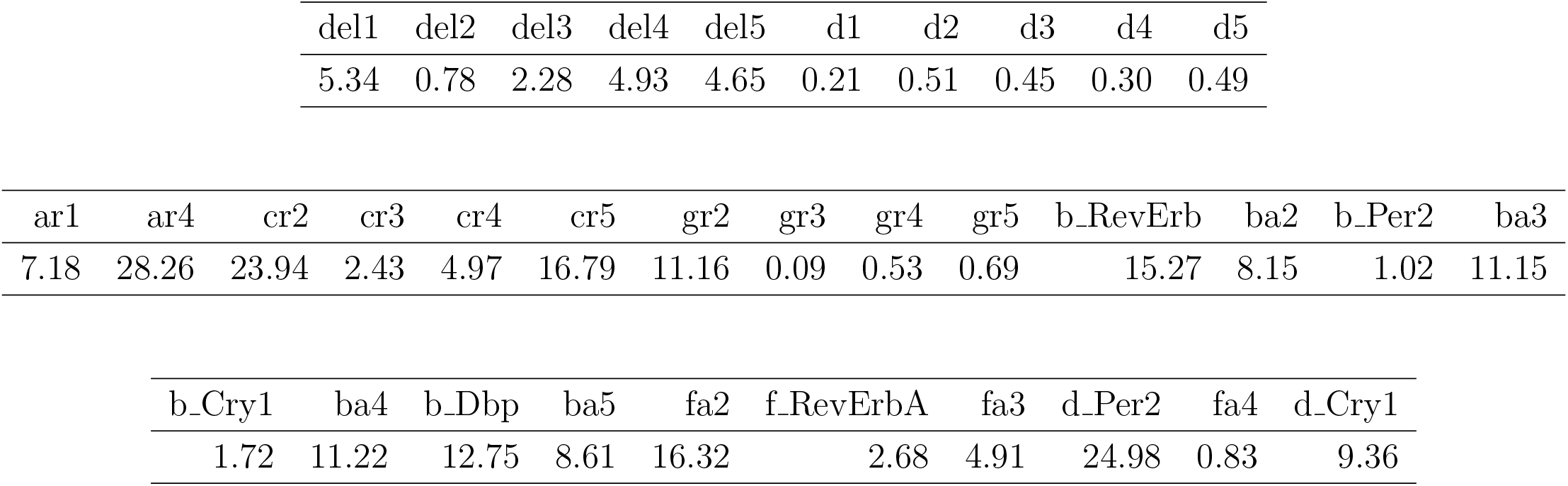

| del1 | del2 | del3 | del4 | del5 | d1 | d2 | d3 | d4 | d5 |
| --- | --- | --- | --- | --- | --- | --- | --- | --- | --- |
| 5.34 | 0.78 | 2.28 | 4.93 | 4.65 | 0.21 | 0.51 | 0.45 | 0.30 | 0.49 |

| ar1 | ar4 | cr2 | cr3 | cr4 | cr5 | gr2 | gr3 | gr4 | gr5 | b_RevErb | ba2 | b_Per2 | ba3 |
| --- | --- | --- | --- | --- | --- | --- | --- | --- | --- | --- | --- | --- | --- |
| 7.18 | 28.26 | 23.94 | 2.43 | 4.97 | 16.79 | 11.16 | 0.09 | 0.53 | 0.69 | 15.27 | 8.15 | 1.02 | 11.15 |

| b_Cry1 | ba4 | b_Dbp | ba5 | fa2 | f_RevErbA | fa3 | d_Per2 | fa4 | d_Cry1 |
| --- | --- | --- | --- | --- | --- | --- | --- | --- | --- |
| 1.72 | 11.22 | 12.75 | 8.61 | 16.32 | 2.68 | 4.91 | 24.98 | 0.83 | 9.36 |

## S3 Scoring parameter sets for global optimization

### 1 Extracting features from experimental data

We fitted harmonic models to experimental data [1] that is publicly available for different tissues with a circadian resolution of 2 h over two days. Using such models, measurements are approximated to yield more reliable estimates of amplitudes and phases.

The fitted models have the form given in Equation S3-1.

Thus, the functions contain harmonics and have curve shapes with wide troughs and narrow peaks. Such a shape is well suited for the measured time series.

The fit parameters a, b and c are obtained by nonlinear regression. Example fits of harmonic models to liver data are shown in Figure S3-1.

**Figure S3-1:**
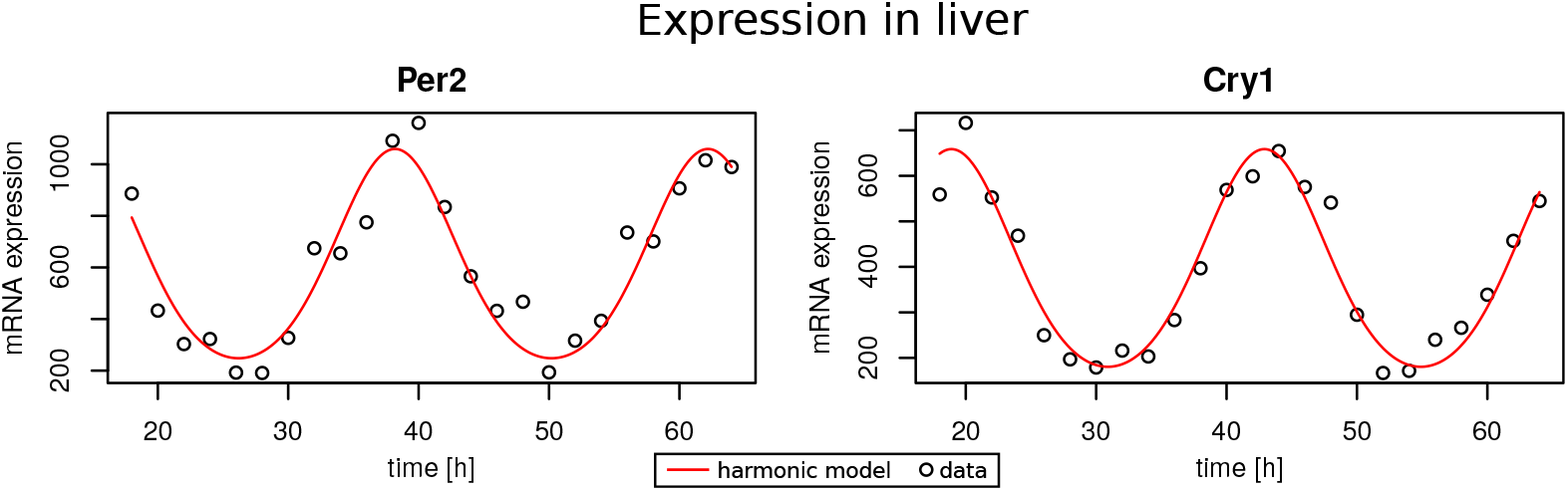
Fitted harmonic models and data points for Per2 and Cry1 in mouse liver.

### 2 The scoring function

The complete scoring function incorporating period, phases and fold changes is given in Equation **S3-2**. Differences between simulated (·_*sim*_) and experimentally measured values (·_*exp*_) are weighted by tolerances (*tol* ·).

As *phase_sim_* and *phase_exp_* relative phase differences to *Bmal1* are used. Thus, there are four phase differences.

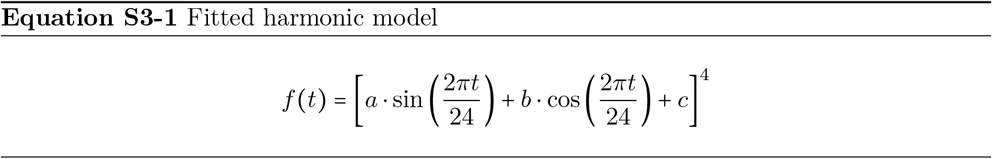

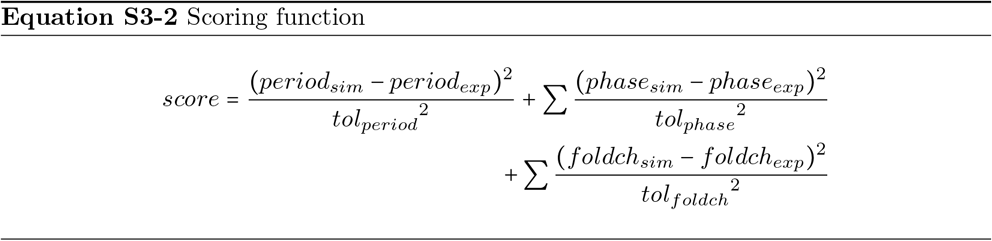

The fold changes are calculated as 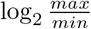. For experimental values we use maxima and minima (peaks and troughs) derived from harmonic fits to the data. In this way, measurement errors of individual points at peaks or troughs are reduced since all points contribute to the fits.

There are 10 terms in total (1 period + 4 phases + 5 amplitudes). If differences between data and fit are equal to the tolerances we get a score of 10. Thus, we consider a score of 10 as a reasonable cutoff.

### 3 Tolerances

We use a tolerance of *tol_period_* = 0.1 h for the period, reflecting typical experimental deviations in mice WT data. For relative phases we compare measurements from different experiments [1–3] and derive a tolerance of *tol_phase_* = 1 h. Figure S3-2 shows that relative phases measured in the three experiments are indeed comparable.

**Figure S3-2:**
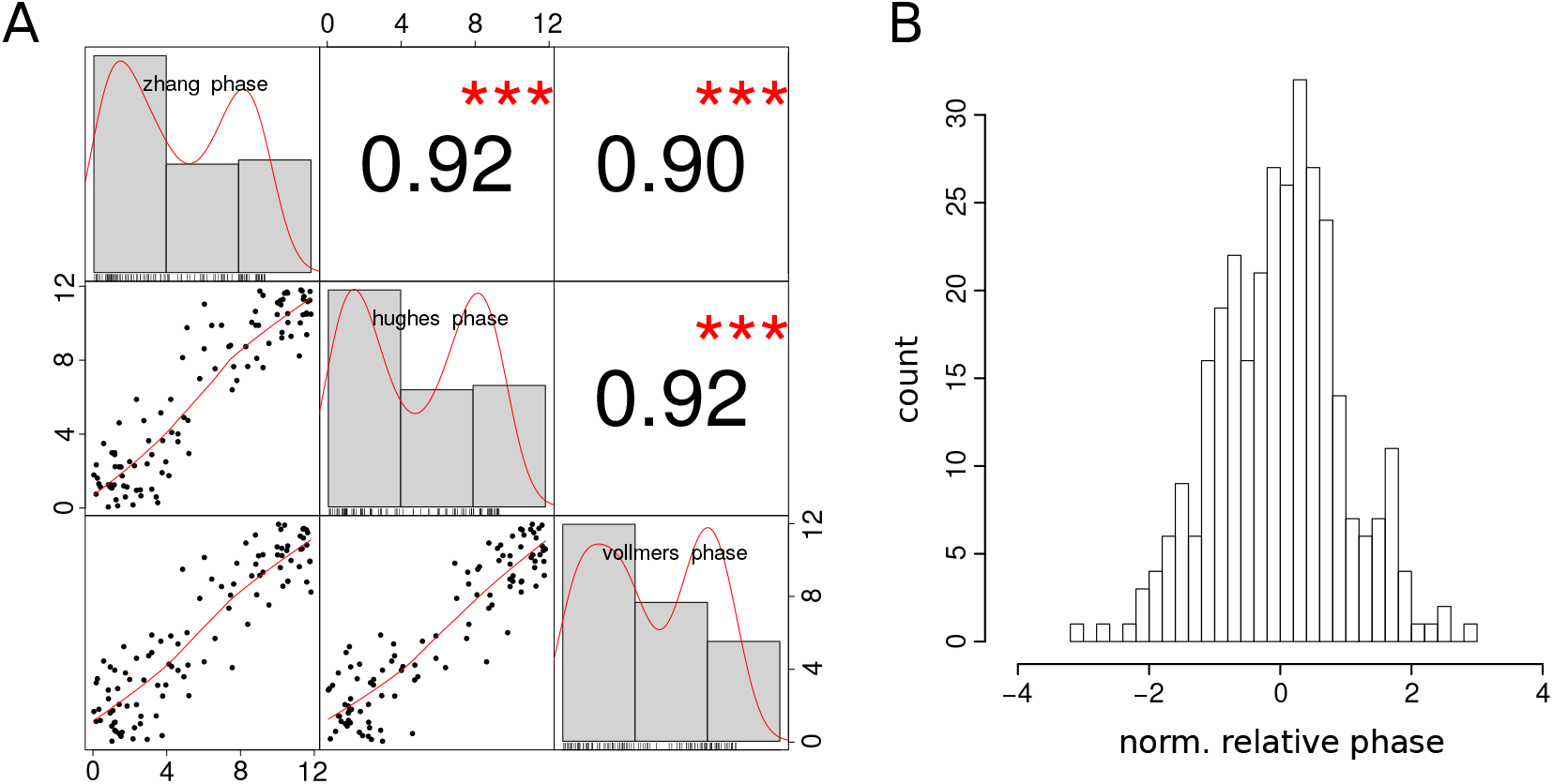
(A) Correlations of relative phase relationships across three experiments showing Pearson correlation coefficients. (B) Histogram of phase differences to the gene-specific mean value. The standard deviation is about 1 h.

Fold changes are less consistent between the different experiments. To define a reasonable tolerance we therefore split the Zhang et al. data set into day 1 and day 2 and compare the deviation between these days. Since fold changes vary strongly between genes, we define five tolerances (one for each gene) based on the median differences between the days: *tol*_*foldch*(*Bmal1*)_ = 0.1, *tol*_*foldch*(*Reverba*)_ = 0.4, *tol*_*foldch*(*per2*)_ = 0.2, *tol*_*foldch*(*Cry1*)_ = 0.1, *tol*_*foldch*(*Dbp*)_ = 0.12.

### Cuts through the fitness landscape

Application of the scoring function to all parameter combinations yields a 35 dimensional landscape with troughs and peaks. Using global optimization with VFO (see Supplement S4) and Particle Swarm Optimization we search troughs with minimal score.

In Figure S3-3 we present a representative selection of cuts through the fitness landscape along one parameter axis. For each depicted parameter, cuts are shown for models fitted to SCN and liver. The parameter values are taken from successful optimization runs and the found minimum is marked in red.

**Figure S3-3:**
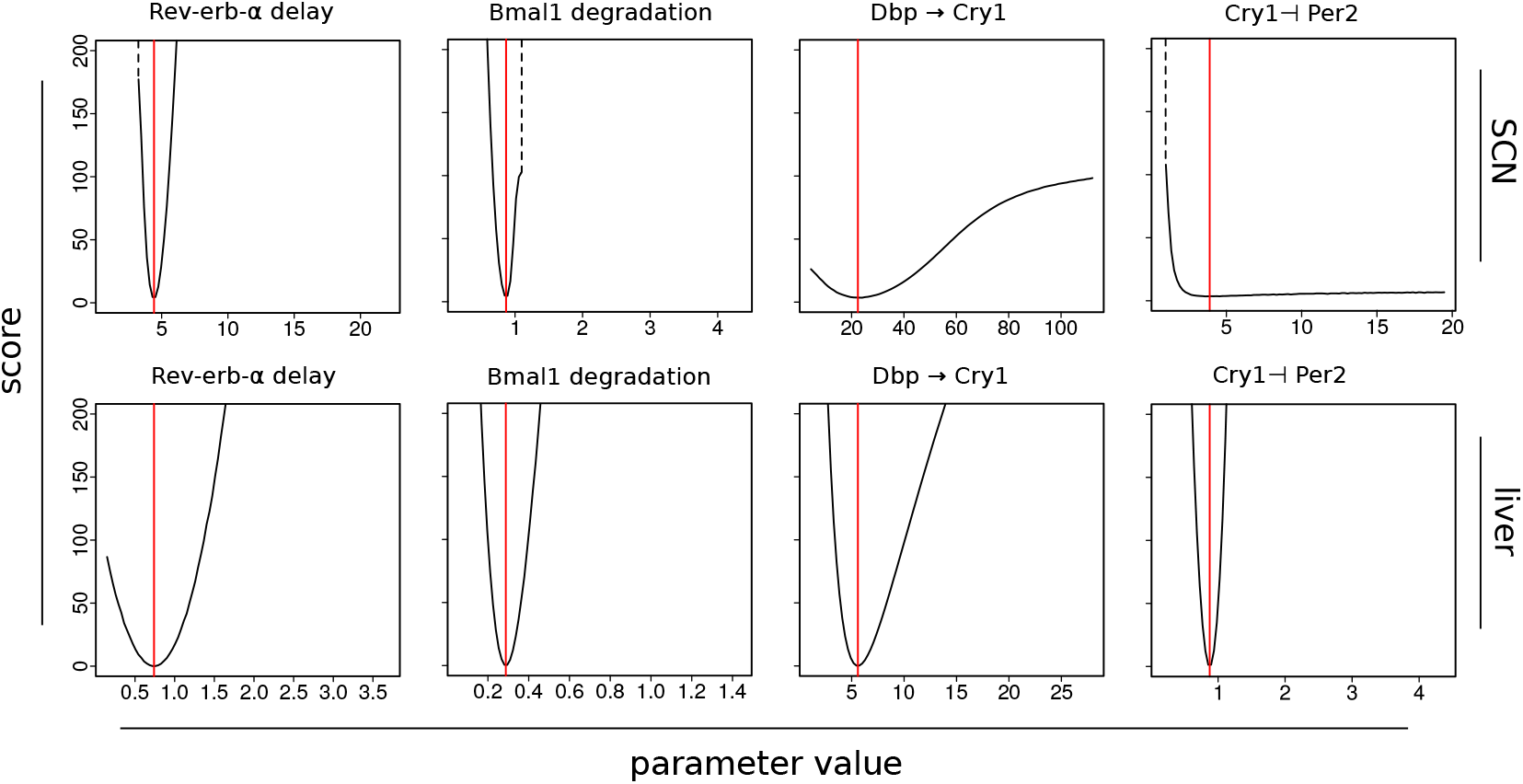
Cuts through the 35 dimensional fitness landscape created by applying the scoring function (Eq. S3-2) to combinations of 34 parameters. In each plot one parameter of an optimal model fit is varied from 0.2 to 5 times the optimal value (red line). Transitions to regions with no oscillation are marked by dotted lines.

This depiction also presents useful information to judge the quality of fits and possibility to identify unique parameter values. Most cuts have a near-parabolic shape and clear trough as in the first 2 columns of Figure S3-3. Only some cuts have minima located in less steep troughs, as for example in row 1, column 3. In fits with less good scores, as for example in row 1, sometimes also a boundary is reached after which rhythms vanish (dotted lines). In just a few cases there is no clear optimum (e.g. last plot in row 1).

Interestingly, *Cry1* ⊣ *Per2*, a regulation that is part of the repressilator, has a clear trough for the fit to liver data which contains a repressilator in its rhythm generating set of loops. However, it shows no discernible optimum for the SCN fit that does not involve this regulation as an essential part of its oscillation generating mechanism.

## S4 Finding initial estimates with Vector Field Optimization

### Fitting the vector field

In our case we have five differential equations, with associated experimental time series. We want to optimize the parameters of these differential equations, such that the solution resembles the time courses. Usually for complex sets of equations this is done with numerical simulations for each parameter set.

With Vector Field Optimization (VFO), however, we aim at fitting the vector field in regions where experimental data is available. Thus, we make use of the situation that time courses exist for all equations.

Fitting is done by plugging in time series values in the right hand side of the differential equations and time series derivative values at the left hand side. This is schematized in Example **S4-1**. Instead of the raw experimental time series, we use harmonic fits (as described in Supplement S3) as well as their analytic derivatives.

**Example S4-1** Plugging in time course values for VFO. In this toy differential equation with an inhibition-, activation- and degradation term the time series values for fits of genes A, B and C are plugged in at the blue places. The time derivative of the time series fit for gene A is plugged in at the green place. Then, for each set of parameter values (red) the difference between the left and right side of the equation is used as a score (sum of squared errors) and the score is minimized using a bounded gradient method. In this way estimates of the unknown parameters (red) are obtained.

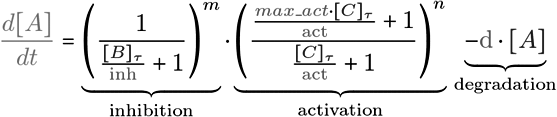

Then, parameters are chosen to minimize the squared differences between both sides of the equation. Since no numeric simulation is required, evaluation of one parameter set is fast. To keep our restrictions on parameter ranges, a bounded gradient method is used for optimizing parameter values.

### Finding initial estimates

We use VFO only for optimizing initial model parameter sets. In principal it is possible to fit a model solely with VFO (if the differential equations can exactly resemble the time courses used for fitting). However, there are shortcomings of this method:

If the fit is not perfect and right hand side and the derivative on the left hand side deviate for one or more equations, it is not guaranteed that the vector field forms asymptotically the desired limit cycle at the respective position. Tests with numeric simulations showed that in some of the optima it indeed forms a limit cycle, in others, however, the gene levels go to a steady state.

We could increase the frequency of found limit cycles by introducing a “transient trick”, requiring vectors in the vicinity of the desired cycle in the vector field to point towards the cycle. However, there were always cases of failure and the resemblance of found limit cycles with time series was overall not sufficient.

Nevertheless, VFO directs parameters to regions in parameter space where time courses and equations roughly match. We therefore tested the use of VFO to compute starting conditions for optimization with a second method. The solutions found with VFO are also used to center initial parameter ranges for the second method around the VFO estimates.

In our implementation we use Particle Swarm Optimization (PSO) as a second method. We find that VFO+PSO yields significantly better scores for liver and SCN than PSO alone (main text Figure 4B), while the scores are comparable in kidney and adrenal gland. Notably, almost no good fits were found in SCN without using VFO. There were no marked differences in the loop distribution when using PSO alone or in combination with VFO.

Further, we use different scoring functions for VFO and PSO that possibly complement each other. While the scoring function used for PSO only optimizes period, phase relations and fold changes, VFO also accounts for the curve shape. VFO finds regions in parameter space, taking the differences between time courses carefully into account, but possibly failing to fit a real limit cycle. Then, PSO is used including numeric simulations to guarantee oscillations with a period of 24 h, correct phase relationships and fold changes of gene expression levels.

### Parameter distribution after VFO

VFO determines starting conditions for PSO and directs the search to certain regions in parameter space. We use the computed result of VFO to initialize 1 of 70 particles used in PSO, while the others are still initialized randomly using latin hypercube sampling. (The score of the VFO particle is mostly very good and pulls other particles into its direction.)

To assess the effect of VFO, we ran 50 optimizations with only VFO on the liver data set and examined the resulting starting estimates (see Fig. S4-1). 37 of 50 of these starting estimates were already oscillating and 7 had scores below 10 using the PSO scoring function. Interestingly, the distributions of many parameters are quite restricted compared to the parameter ranges, suggesting specific optimal values.

**Figure 8:**
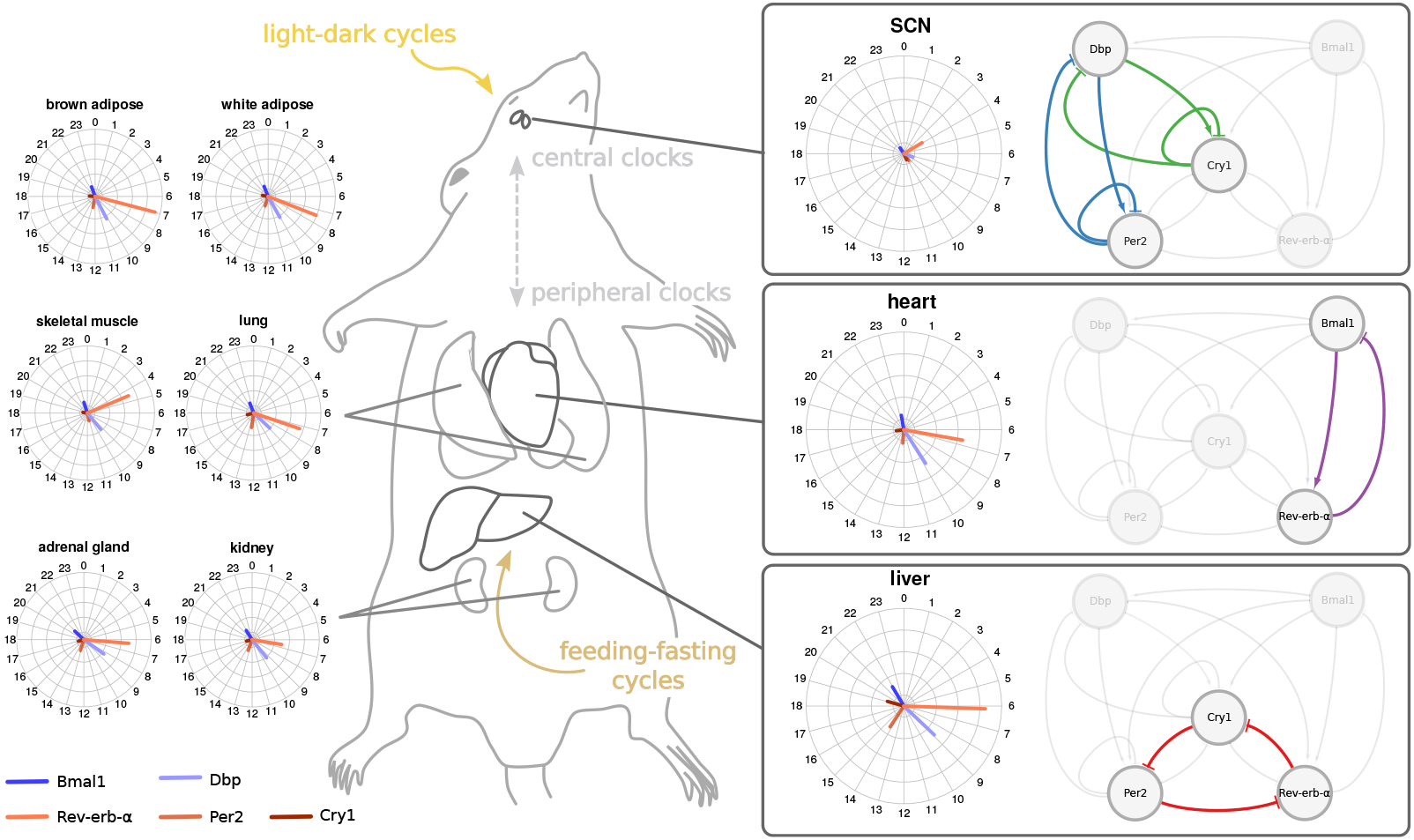
Differences between gene expression in mouse tissues and associated feedback loops found in model fits. Shown are three representative feedback loops for tissues as characteristic examples.

For example the delays of *Bmal1* (del1) and *Rev-erb-α* (del2) are about 6 and 1 respectively. This relation is consistent with the time gap between peaks of those genes and with the *Bmal1-Rev-erb-α* feedback loop in particular. Further the kinetic parameters corresponding to this loop (ar1, b_RevErb and ba2) are also contained in narrow ranges. Indeed, our clamping analysis revealed that *Bmal1-Rev-erb-α* loops are prevalent.

The emergence of such specific parameter patterns suggests that it is reasonable to pre-emphasize the search for optima with VFO. Improved scores and pre-selection of parameter sets indicate, that VFO is well suited to improve optimization.

**Figure S4-1:**
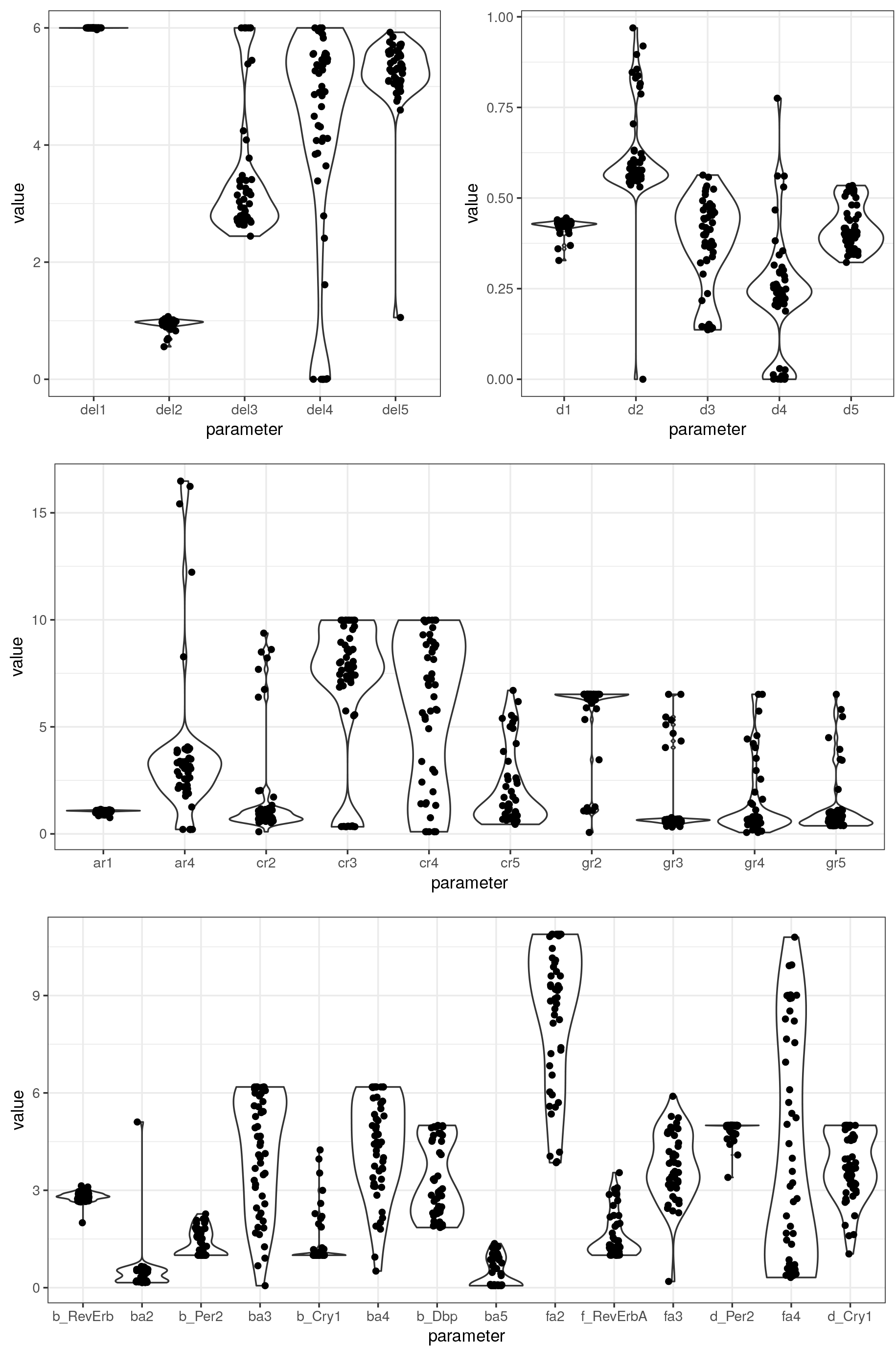
Parameters after VFO. The resulting parameter distributions deviate clearly from equidistributions in the alowed parameter ranges such as [0,6] for delays and [0,1] for degradation rates.

## S5 Identification of essential feedback loops

### Clamping regulations

Given a fitted parameter set, we analyze which loops in the network are essential for the generation of rhythms. To this end we employ our clamping strategy that was published earlier [1]: By setting combinations of regulatory links in the network graph to a constant value we systematically test which links are necessary for oscillations.

Let us assume that a regulatory link is part of an essential loop. Then rhythms vanish if we set this regulation to a constant. We choose the mean values of oscillations in the default state as a constant to preserve the basal effect exerted by the clamped regulation.

Any combination of regulations can be clamped by setting the respective parts in the differential equations constant.

### Combinatorial exploration of the loop structure

To avoid excessive computation times, we do not test all combinations of regulations (which are 2^17^ = 131072). Instead we resort to a targeted clamping strategy.

It is known that a negative feedback loop is necessary to generate oscillations [2]. Therefore, we list and test negative feedback loops in the network specifically.

We start by testing each single loop individually: A negative feedback loop is termed essential, if separate clamping of all regulations involved in the loop (i.e. clamping one regulation at a time) leads to disruption of rhythmicity. (Note, that in theory there might be a case where all edges of a negative feedback loop are shared by other loops, making it impossible to attribute the effect of clamping in that way, but this is not the case in our network.)

After testing single loops we proceed by testing combinations of loops. This is of interest, since loops could mutually compensate for each other. Then clamping just a single loop does not stop rhythms, because the other loop still generates oscillations and vice versa. In analogy to the single loop case we define a combination of loops as essential, if separate clamping of all pairwise combinations of regulations from the two loops leads to disruption of circadian rhythmicity. An example is given in Figure S5-1.

**Figure S5-1:**
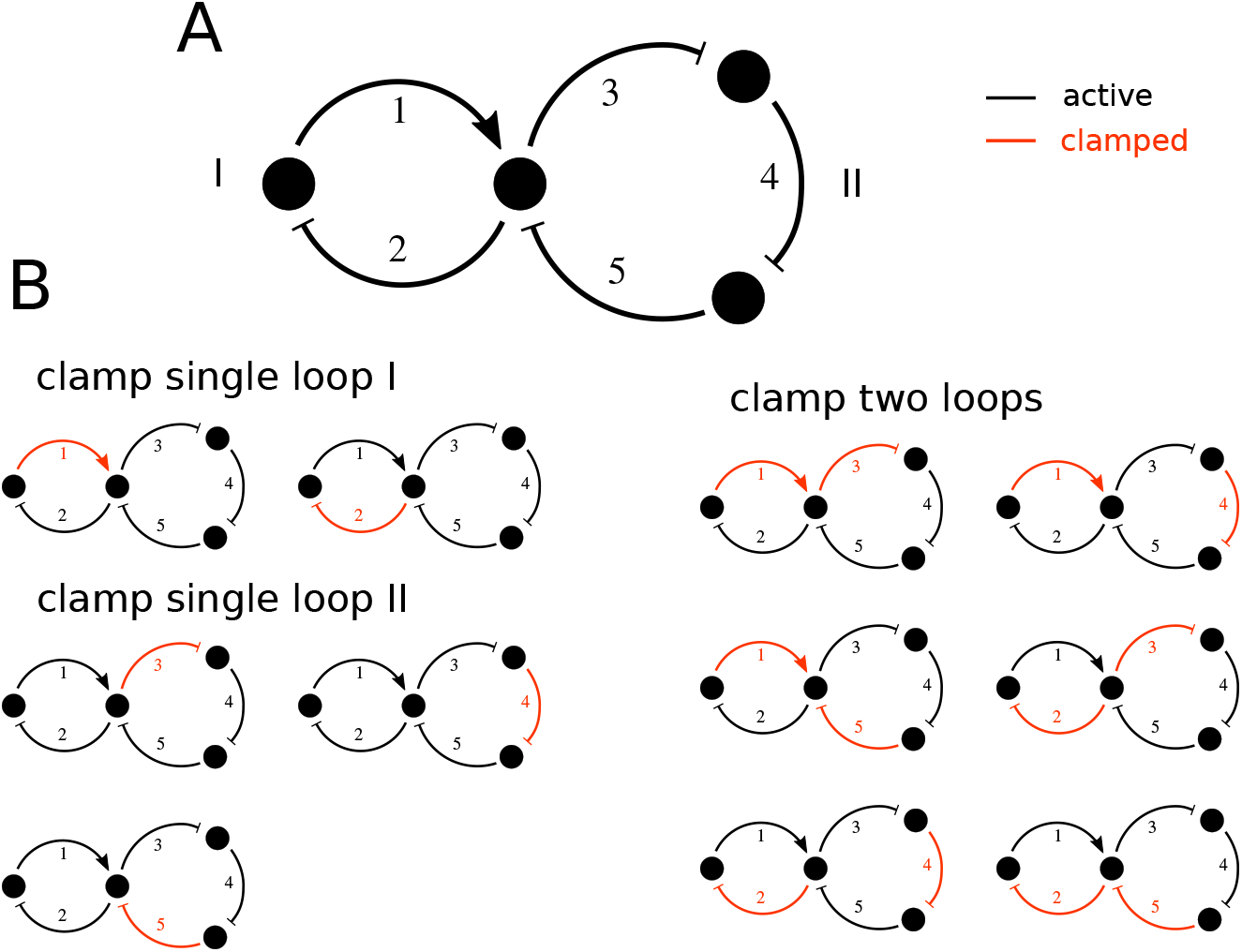
Example for targeted clamping of loops. (A) A toy network with two negative feedback loops ({1,2} and {3,4,5}). Assume that the model represented by this network is rhythmic, then both loops are candidates for the core mechanism generating these rhythms. The mechanism could involve loop I, loop II or both. (B) On the left: Clamping of single loops. To test whether e.g. loop II is essential, the model is simulated three times, with regulations 3, 4, and 5 being clamped in one simulation each. If all three simulations show disruption of rhythmicity loop II is regarded as essential. On the right: Clamping the combination of loop I and II. If all six simulations show disruption of rhythmicity the combination of loop I+II is regarded as essential. If both single loops I and II are essential, we denote this as I ⋁ II, i. e. interrupting any loop stops oscillations. If the combination of loop I and II is essential, we denote it as I ⋀ II, i.e. interrupting both loops simultaneously stops oscillations.

If two single loops were essential, then the combination of both loops is also essential. Therefore we do not need to test the combinations of loops that have already been identified as single essential loops. If a combination of loops is essential, but not the single loops, these loops mutually compensate for loss of one of the loops.

An overview over the procedure is shown as pseudocode in Algorithm 1.

**Algorithm 1.**
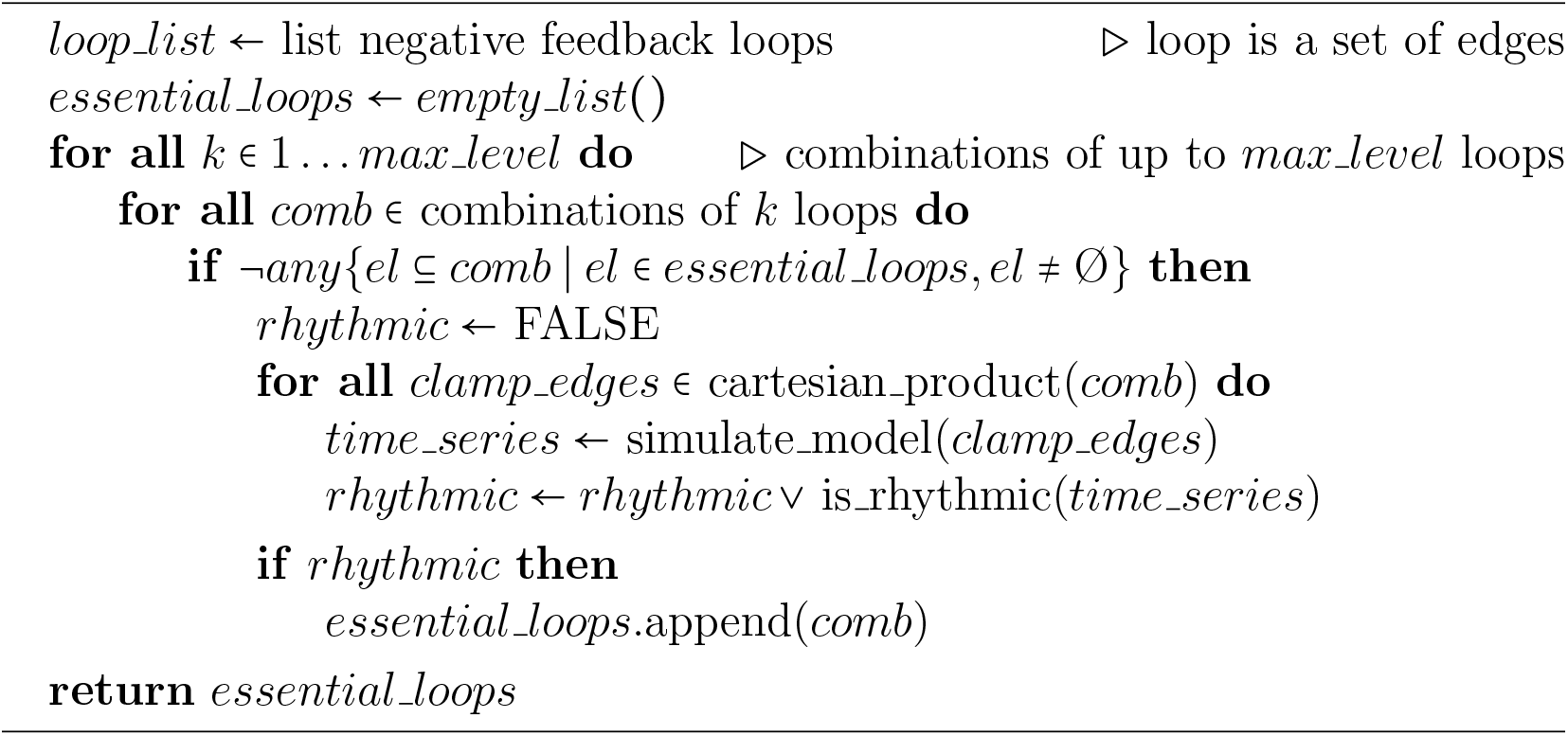
targeted clamping.

### Output notation and oscillators

The algorithm returns, a list of feedback loops needed to be clamped to disrupt rhythmicity. We can denote this in the form of a logic function. For the example in Figure S5-1A, we could have for instance *I* ⋁ *II* (see Figure caption).

Assuming a more complex network with 5 negative feedback loops, we could also have:

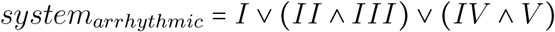

with *I … V* being 1 if the corresponding loop is clamped and 0 otherwise. It means that clamping loop *I* disrupts rhythms and also clamping the combination of loops *II* and *III* as well as loops *IV* and *V*, while clamping loop *II* or *V* alone does not.

This condensed description in Disjunctive Normal Form (DNF) is useful to see the different options of how to disrupt rhythms with a minimal amount of clamped regulations.

If we want to see on the other hand, which minimal amount of loops is necessary to generate rhythms, we can transform the function (by negating, applying De Morgans law and converting back to DNF):

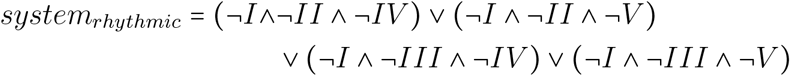

¬*I* … ¬*V* are 1 if the corresponding loop is active and 0 otherwise. The expression now describes, which feedback loops need to be active in order for the system to be rhythmic.

In other words: each combination of loops inside the brackets (e.g. *I, II* and *IV*) constitutes a minimal oscillator. Thus, all brackets contain minimal sets of loops that can generate oscillations in synergy.

We identify minimal oscillators for all logic functions and verify them in separate simulations. Thus, we are able to find necessary and sufficient conditions for rhythms in an automated way in all fitted models.

### Loop frequencies in different tissues

We used our clamping analysis to find essential loops in all fitted models. Interestingly a large variety of loops and oscillators was found (see main text). Figure S5-2 shows the relative frequencies of loops found in 10 tissues. To keep the legend simple, essential loops were counted if they were present in the logic function determined by our method (see above), regardless of their position in this function.

---

**Figure S5-2:**
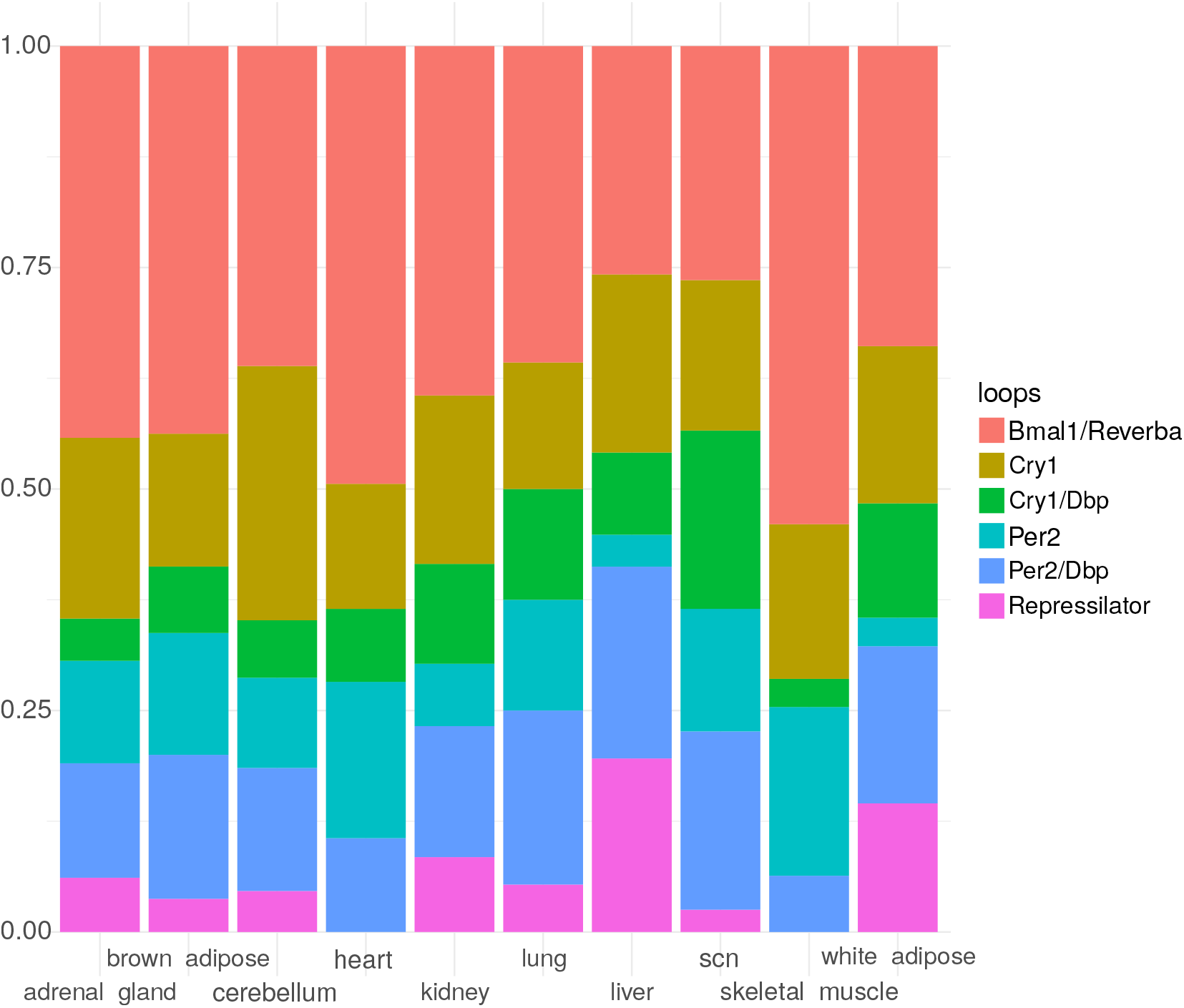
Relative loop frequencies in 10 tissues. In skeletal muscle and heart no repressilator was found, but they have a large fraction of models with essential *Bmal1-Rev-erb-α* loop. In SCN the fraction of combinations with *Per2* and *Cry1* loops is particularly large and most repressilators were found in liver.

